# Biomolecular condensate architecture of an autophagic cargo at molecular resolution in situ

**DOI:** 10.64898/2026.01.07.698105

**Authors:** Emily Boyle, Sergio Cruz-León, Javier Lizarrondo, Johann Brenner, Hector Mancilla, Jan F. M. Stuke, Jorge Jimenez-Niebla, Sigrid Milles, Sonja Welsch, Carola Hunte, Claudine Kraft, Gerhard Hummer, Florian Wilfling

## Abstract

Biomolecular condensates organise cellular biochemistry, yet their molecular architecture in situ remains poorly understood. During selective autophagy, macromolecules frequently accumulate into biomolecular condensates, forming discrete entities for autophagic engulfment and degradation - ideal systems for structural analysis. We employed in situ cryo-electron tomography to determine the near-atomic resolution structure of Aminopeptidase 1 condensates within cells. These condensates form densely packed, spherical assemblies with amorphous organisation and liquid-like properties, elucidating the requirements of a selective autophagic cargo for exclusive targeting. Structural analysis and multiscale simulations reveal that the short, transient α-helical structures in the disordered N-terminus of Aminopeptidase 1 enable site-specific, coiled-coil-like interactions required for condensate formation and properties. A single point mutation that increases α-helical propensity directly modulates condensate viscosity and dynamics from a liquid-like to a glass-like state, while preserving local molecular packing. Our results demonstrate that disordered regions encode both specificity and material properties through transiently structured motifs, linking sequence specificity to phase behaviour in cells and expanding the molecular logic of phase separation in cells.

## Main

Membraneless organelles are dense assemblies of biomolecules, often arising through processes that resemble phase transitions^1–3^. These cellular biomolecular condensates create specialized subcellular microenvironments and thereby mediate various processes, including gene expression, nuclear transport, signalling and stress responses^4,5^. Biomolecular condensates can exhibit liquid-, gel-, or solid-like states, ranging from low-viscosity liquids to viscoelastic gels, and frequently form glassy or fibrillar amyloid networks under ageing or pathological conditions^6^. The material properties and composition of biomolecular condensates determine their cellular function, and dysregulation of their formation has been associated with cancer, neurodegenerative diseases, and ageing^7^. Mechanistic insights into their intermolecular interactions and the resulting supramolecular organisation are essential to unravel their role in disease and to inform targeted therapeutic strategies.

Although in vitro studies using recombinant proteins have provided foundational principles of condensate formation^8^, elucidating their organisation in the crowded and heterogeneous cellular environment remains a major challenge. Cryo-electron tomography (cryo-ET) enables in situ visualization of macromolecular assemblies at molecular resolution; however, the inherent structural disorder and compositional complexity of many condensates have so far limited high-resolution analyses in cells^9–12^.

To address these limitations, we turned to functional condensates formed by aminopeptidase-1 (Ape1), a vacuolar enzyme in *Saccharomyces cerevisiae (S.c.)*, that is targeted by selective autophagy. Unlike many condensates, Ape1 condensates have compositional simplicity and relatively low structural disorder, making them particularly tractable for in situ cryo-ET. Briefly, Ape1 assembles into tetrahedral homododecamers that form condensates within the cytoplasm^13,14^. These condensates exhibit liquid-like properties and are stabilized by interactions between N-terminal propeptides protruding from the dodecamers (Fig. 1a)^15,16^. Remarkably, the otherwise disordered propeptides adopt a helical structure in in vitro experiments, forming coiled-coil-like interfaces with neighbouring propeptides^14^. Ape1 condensates are then delivered to the vacuole by a canonical selective autophagy pathway, the cytoplasm-to-vacuole targeting (Cvt) pathway^17^. The propeptide is cleaved in the vacuole, resulting in condensate disassembly and dispersal of the mature, active form of Ape1 (mApe1). Here, we use in situ cryo-ET to reveal the spatial organization of this protein condensate at unprecedented resolution and, together with molecular simulations, to relate the distinct molecular features of Ape1 to condensate material properties in the native environment, and in turn to biological function.

**Figure 1:**
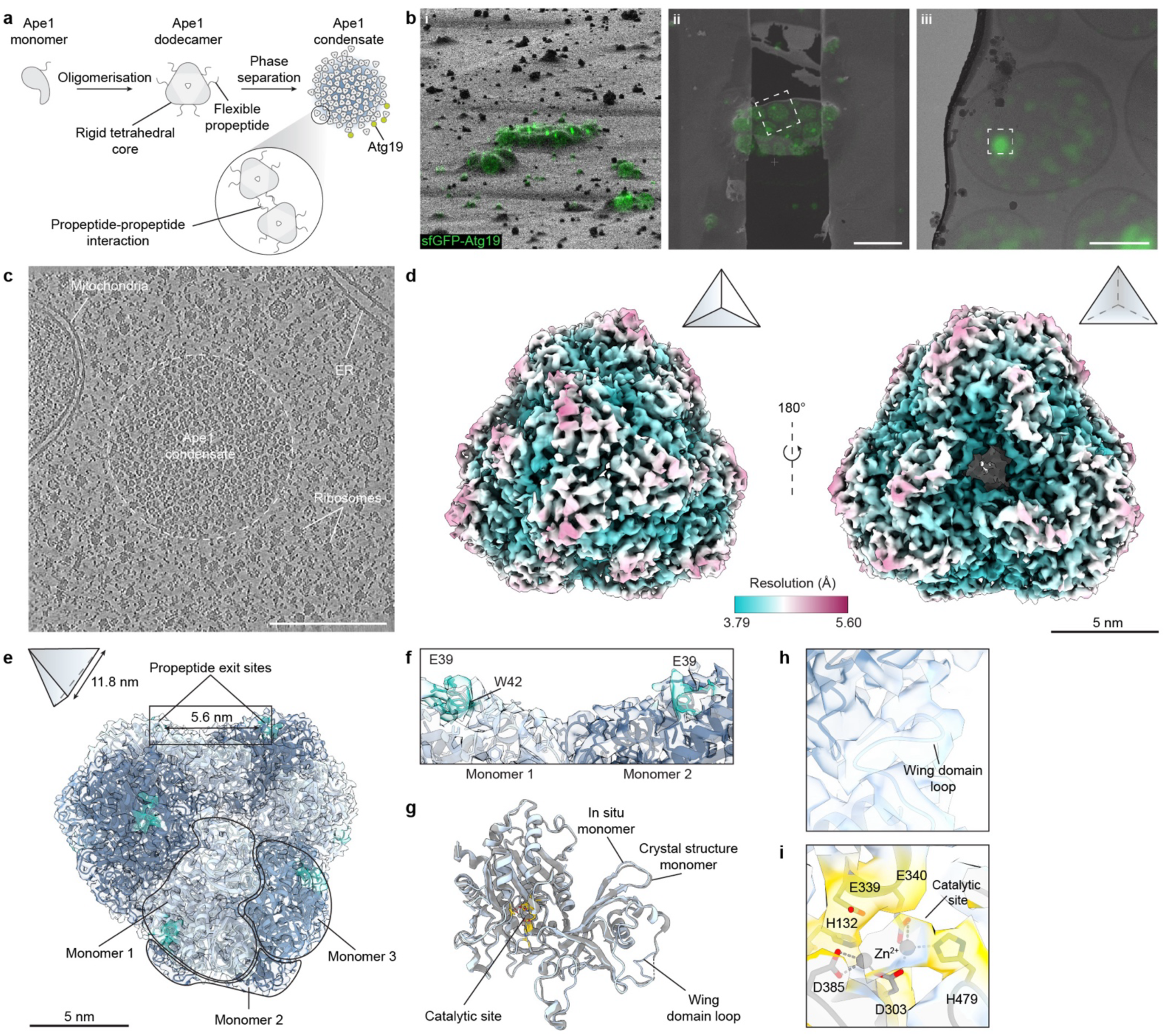
In situ structure determination of Ape1 dodecamers by correlative cryo-ET. **a)** Schematic of Ape1 condensate formation in *S. cerevisiae*. **b)** Example of correlative cryo-ET pipeline to target Ape1 condensates inside cells. Maximum intensity projection of a sfGFP-Atg19 cryo-fluorescence stack is overlaid on a focused ion beam image of the milling target (i), a scanning electron microscope image of the milled lamella (ii), and a transmission electron microscope image of the lamella (iii) at the region indicated by the white box in (ii). Scale bars = 5 µm (ii) and 2 µm (iii). **c)** Central slice of a denoised tomogram from the region indicated by the white box in Fig. 1b)iii). Scale bar = 250 nm. **d)** In situ electron density map of the Ape1 dodecamer coloured by local resolution. Small tetrahedra indicate dodecamer orientation. Scale bar = 5 nm. **e)** Ape1 dodecamer model built into the map. Monomers are differentially coloured around the 3-fold symmetry axis and outlined at one vertex. Residues 38-45 are coloured in cyan. Small tetrahedron indicates dodecamer orientation. Scale bar = 5 nm. **f)** Edge of the Ape1 tetrahedron indicated by the black box in Fig. 1e). Resolved residues 38-45 of the propeptide are coloured in cyan. **g)** In situ Ape1 monomer (light blue with catalytic residues highlighted in yellow) overlaid with the Ape1 L11S crystal structure (PDB: 5JH9, grey)^14^. **h)** View of the wing domain loop backbone modelled into the in situ map. **i)** Fitting of catalytic residues from the Ape1 L11S crystal structure (grey) into the in situ map^14^. Catalytic residue density is shown in yellow.

### Near atomic resolution structure determination enables the molecular study of Ape1 condensates in situ

To study the organisation of Ape1 condensates at molecular resolution in situ, we employed a 3D-correlative focused ion beam (FIB)-milling and cryo-ET workflow (Fig. 1b)^18^. While fluorescent labelling of the protein of interest is a common detection strategy, labelling of Ape1 is likely to affect its properties, including intermolecular interactions within its higher-order assemblies, as observed for other phase-separating proteins^19–21^. Therefore, to preserve the native state of the Ape1 condensate, we instead labelled the selective autophagy receptor Atg19, which is recruited to the surface of Ape1 condensates (Fig. 1a and Fig. S1a) and thereby serves as an ideal positional marker for in situ cryo-ET imaging without affecting Ape1 condensate formation^22,23^.

To investigate Ape1 condensate formation during autophagy under nutrient-rich conditions, we treated cells with the mTOR inhibitor rapamycin (Fig. S1a). In addition, Ape1 was overexpressed from a CuSO4-inducible promoter to increase Ape1 condensate size, thereby improving the success rate of targeting by correlative cryo-ET while still permitting phagophore formation and recruitment of associated machinery^24^. Under these conditions, Atg19 tagged with superfolder GFP (sfGFP-Atg19) was visible as either puncta or rings inside cells, reflecting the heterogeneity in Ape1 condensate size regardless of rapamycin treatment (Fig. S1b). Targeting sfGFP-positive structures resulted in a total of 30 tomograms containing distinct accumulations of tetrahedral-shaped protein densities separated from the bulk cytosol, reminiscent of the Ape1 condensate (Fig. 1b and 1c, Fig. S1c and S1d and Supplementary Video S1)^25^. We found that 66% of tomograms showed Ape1 condensates near the vacuole, and 30% had a phagophore surrounding a portion of the condensate (Fig. S1e). This finding is consistent with observations that phagophore initiation during selective autophagy occurs at the Ape1 condensate surface and involves interactions between phagophore initiation machinery proteins and the vacuolar membrane protein Vac8^18,25,26^. In addition, parts of the endoplasmic reticulum were observed in 66% of our tomograms, in accordance with its role in contributing lipids to phagophore membrane growth (Fig. S1e)^27^.

To dissect Ape1 condensate organisation in situ, we performed template matching combined with subtomogram averaging (STA). After obtaining an initial reference using the *S. pombe (S.p.)* Ape4 cryo-EM structure as a template^28^, the resulting map was used for high-confidence 3D template matching (hcTM) on the whole dataset using GAPSTOP^TM29^. hcTM provided strong cross-correlation peaks within the accumulations of tetrahedral densities in our reconstructions (Fig. S2). After 3D classification and refinement imposing tetrahedral symmetry, the final reconstruction of Ape1 reached a global resolution of 3.84 Å (Fig. 1d and Fig. S2).

At this resolution, we were able to dock the previously determined crystal structure of the Ape1 L11S mutant^14^ and generate an in situ atomic model of Ape1 (Fig. 1e and Fig. S3a). Consistent with existing crystal structures, our in situ structure of the Ape1 dodecamer adopts a tetrahedral architecture, with three Ape1 monomers positioned at each vertex (Fig. 1e)^14,30^. In this arrangement, propeptides from two Ape1 monomers are located at each of the edges of the tetrahedron, with an edge length of 11.8 ± 0.1 nm and a distance between propeptides of 5.6 ± 0.4 nm (Fig. 1e and 1f). The N-terminal portion of the propeptide (residues 1-37) was unresolved, with the density allowing us to confidently build a model only from residue 38 onwards (Fig. 1f and Fig. S3b). In existing cryo-EM and crystal structures, the propeptide also remains largely unresolved due to its flexibility, as demonstrated by previous NMR chemical shift analysis^14^. However, the resolution of our map allowed us to assign the positions for the backbone atoms of a flexible loop in the wing domain of Ape1 (Pro 234 to Pro 240), which could not be modelled in the previous crystal structures (Fig. 1g and 1h)^14,30^. All catalytic residues located towards the core of the dodecamer are in good agreement with the Ape1 L11S crystal structure (Fig. 1i and Fig. S3c). Remarkably, density was also observed at the position corresponding to one of the catalytic zinc ions, characteristic of the M18 family of metalloproteases to which Ape1 belongs (Fig. 1i and Fig. S3c)^14^. Finally, a model of the structurally similar *S.c.* Ape4 dodecamer fit poorly into our map, demonstrating the absence of template bias in our Ape1 structure (Fig. S3d and S3e). Our in situ structure of Ape1 provides compelling evidence that the targeted Ape1 condensates consist of densely packed dodecamers, and demonstrates that 3D-correlative cryo-ET enables high-resolution structure determination of a ∼630 kDa protein within a crowded condensate environment inside cells.

### Ape1 condensates form dense spherical assemblies with an amorphous organisation

Visual inspection of Ape1 condensates revealed a predominantly spherical morphology (Fig. 2a and 2b), with sphere-fitting analysis yielding a median diameter of ∼466 nm (Fig. 2c and Fig. S4a-c), consistent with previous reports of Ape1 condensate size under mild Ape1 overexpression^24^. This sphericity suggests a surface tension-driven assembly of the Ape1 dodecamers in line with previous in vitro work^16^. To quantitatively assess the packing of Ape1 dodecamers within the condensate, we calculated the radial distribution function (RDF)^31^, which measures the probability of finding another Ape1 dodecamer at a distance *r* from a reference particle relative to a random distribution (Fig. 2d). The RDFs were normalised to the volume of a concave hull occupied by Ape1 dodecamers to account for shape irregularities (Fig. S4d). The RDFs displayed three damped oscillations with a periodicity of approximately 13 to 15 nm, matching the circumsphere diameter of an Ape1 dodecamer. Oscillations were followed by signal decay beyond the third peak indicating only short-range order as expected for a dense, liquid-like assembly (Fig. 2d and Fig. S4e). Furthermore, Fourier transforms (FFTs) of the Ape1 particle coordinates showed a concentric ring at ∼1/13 nm rather than discrete Bragg peaks of a crystal lattice, further indicating an amorphous, liquid-like organisation (Fig. S4f). Given that the physicochemical properties of proteins in condensates can regulate client recruitment^32^, we reasoned that the high density of Ape1 dodecamers would exclude random cytosolic proteins from entering the condensate and being aberrantly delivered to the vacuole. Indeed, cytosolically overexpressed mScarlet, which has a molecular weight of ∼25 kDa, was largely excluded from the condensate interior (Fig. 2e and 2f). Together, these analyses establish, at the molecular level, that Ape1 condensates in cells are densely packed amorphous assemblies with a liquid-like character, comparable to other condensates characterised by cryo-ET, such as the RuBisCO pyrenoid^10,11,33^.

**Figure 2:**
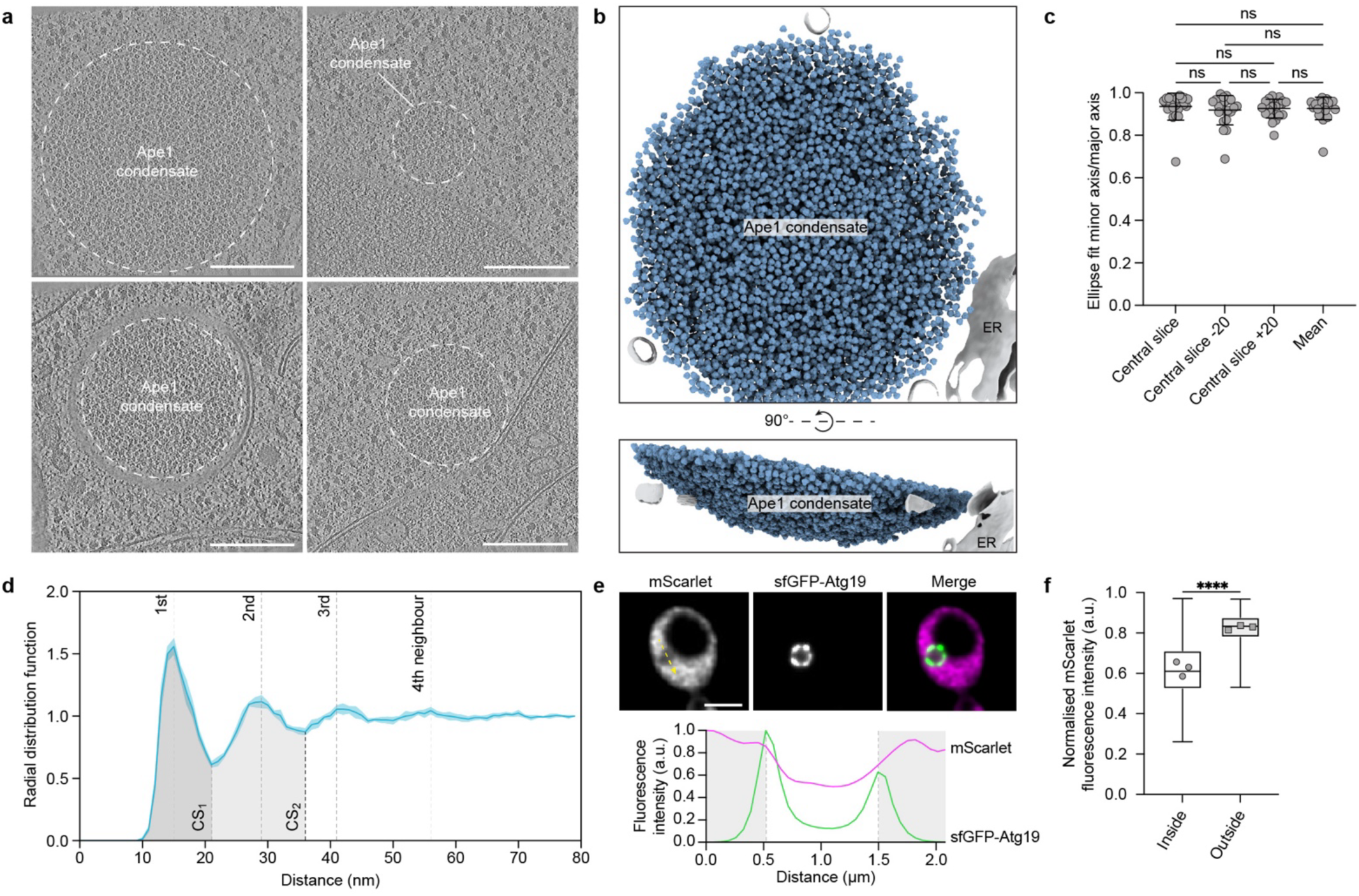
Ape1 condensates are dense, liquid-like assemblies of Ape1 dodecamers. **a)** Central slices of representative denoised tomograms containing Ape1 condensates. Scale bars = 250 nm. **b)** Segmentation of a representative tomogram indicating Ape1 particle positions and surrounding ER membrane. **c)** Ratio of minor and major axes of ellipse fits of segmentations of single slices through 23 Ape1 condensates. Error bars represent the mean ± standard deviation. Non-significance (ns, multiplicity adjusted P-value > 0.05) calculated by one-way ANOVA with Tukey multiple comparisons test. **d)** RDF of Ape1 condensates. Solid blue line is the average profile calculated from 4 tomograms and shaded area indicates the standard deviation. Dashed lines show successive neighbour shells and grey shaded regions delineate the first and second coordination spheres (CS1 and CS2). **e)** Top: Single slices of representative deconvolved fluorescence microscopy images showing exclusion of cytosolically expressed mScarlet from the Ape1 condensate surrounded by sfGFP-Atg19. Scale bar = 3 µm. Bottom: Line profile showing measured fluorescence intensity (normalised to the maximum intensity for each channel) along the yellow arrow indicated in the mScarlet image. **f)** Mean normalised fluorescence intensity of mScarlet inside and outside of sfGFP-Atg19 signal maxima along line profiles through Ape1 condensates from 3 biological replicates. >150 condensates were measured per replicate. Box plot shows the 25^th^ and 75^th^ percentiles with whiskers extending to the maximum and minimum plotted values. Horizontal line shows the median of all measured condensates and individual data points show mean fluorescence intensity per biological replicate. **** P < 0.0001 calculated by paired, two-tailed t-test.

### A single point-mutation reshapes Ape1 condensates at the mesoscale while preserving local structure

While various mutations in the propeptide inhibit Ape1 condensation^14,34–36^, a proline-to-leucine substitution (P22L) alters the properties of the Ape1 condensate from liquid-like towards a more solid state and abolishes autophagic turnover by the Cvt pathway^13,16^. This mutant illustrates how a single substitution can drive a phase transition reminiscent of pathological protein aggregation, and that such physical changes directly impair function^13,16^. Nevertheless, the underlying molecular mechanism remained unknown.

To characterise the impact of the P22L substitution on condensate structure and organisation, we applied our correlative cryo-ET workflow to cells overexpressing Ape1 P22L in the presence or absence of endogenous wild-type (WT) Ape1 (Fig. S5a). Compared to Ape1 WT, Ape1 P22L showed diminished processing into mApe1, consistent with defective delivery of Ape1 P22L condensates to the vacuole (Fig. S5b). Nevertheless, Ape1 WT and Ape1 P22L displayed similar frequencies of condensate formation (Fig. S5c and S5d) and maintained Atg19 recruitment to the condensate surface (Fig. S5e and S5f). As strains with or without endogenous Ape1 WT showed comparable condensate morphology and organisation, they were pooled into a single dataset, referred to subsequently as P22L.

We observed dramatic differences in the P22L and WT condensates at the mesoscale. P22L condensates appeared less spherical, forming various irregular shapes. Strikingly, we also observed holes inside the P22L condensates that were devoid of Ape1 dodecamers (Fig. 3a and 3b and Fig. S5a). Although the content of these inclusions was unclear, cytosolic ribosomes were notably absent (Fig. S5g). We also observed a drastic decrease in the association of Ape1 condensates with phagophore membranes, dropping from 30% of tomograms showing Ape1 WT condensates in the process of phagophore engulfment to 11% for P22L condensates, consistent with impaired autophagic trafficking (Fig. S5h)^13,16^.

**Figure 3:**
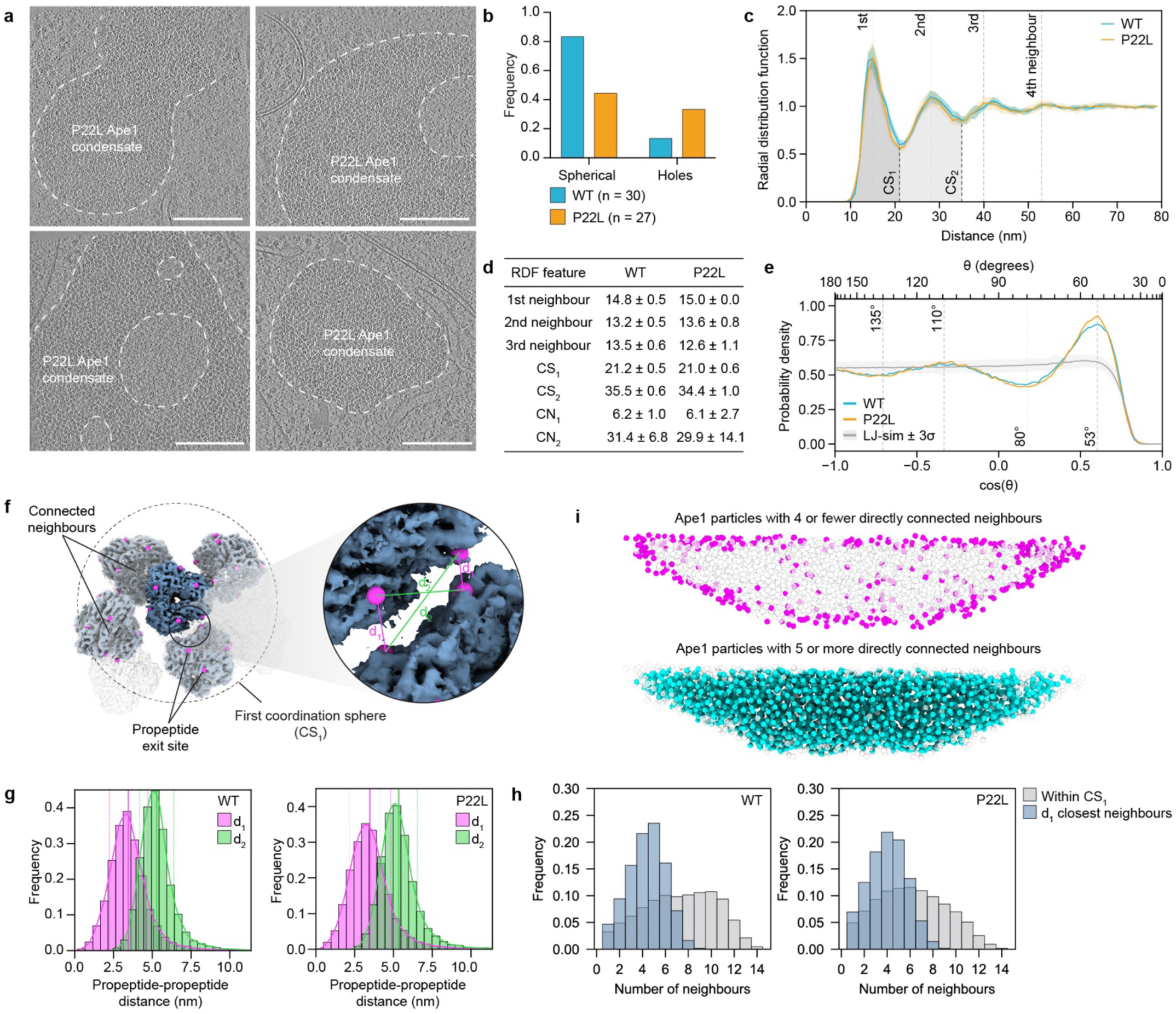
**P22L Ape1 condensate structure is altered at the mesoscale with no change in local organisation of Ape1 dodecamers**. **a)** Central slices of denoised tomograms containing P22L Ape1 condensates. Scale bars = 250 nm. **b)** Frequency of tomograms containing WT and P22L Ape1 condensates with observed morphological features. **c)** RDFs of WT (blue) and P22L (orange) condensates. Solid lines are the average profiles from n = 4 WT and n = 7 P22L tomograms and shaded area indicates the standard deviation. Dashed lines show successive neighbour shells and grey shaded regions delineate CS1 and CS2. **d)** Position (nm) of RDF features for WT and P22L Ape1 condensates, with coordination numbers CN1 and CN2 for coordination spheres CS1 and CS2. **e)** Probability density of cos(θ) for all Ape1 triplets within CS1 of the central particle, where θ is the triplet angle at the centre. The grey line indicates the probability density for randomly distributed spheres from a LJ-simulation of fluid argon with three standard deviations shown as the shaded area. **f)** Central Ape1 dodecamer and its neighbours within CS1. Connected neighbours, as determined by the minimum propeptide-propeptide distances, are shown in blue. Propeptide exit sites (residue 40) are indicated as magenta spheres. The zoom-in shows a single interface between two dodecamers, indicating the closest propeptide-propeptide distance (d1) and the second closest propeptide-propeptide distance (d2) for each propeptide exit site. **g)** Frequency of the closest (d1) and second-closest propeptide-propeptide distances calculated for all Ape1 dodecamers across 4 WT and 7 P22L Ape1 condensates. **h)** Frequency of number of neighbours for each Ape1 dodecamer defined by all particles within CS1 or the minimum propeptide-propeptide distance. **i)** Cross-section through the xz plane of the Ape1 condensate shown in Fig. 2b). Ape1 dodecamers are represented as spheres coloured according to the number of directly connected neighbours as defined in Fig. 3f).

By contrast, at the microscale, the P22L and WT Ape1 condensates were nearly indistinguishable. The overall structure of the P22L Ape1 dodecamer, determined by STA at a global resolution of 5.6 Å showed no observable differences from WT, indicating that the propeptide mutation does not cause structural rearrangements within the dodecamer (Fig. S6 and S7). Moreover, the spatial organisation of P22L and WT condensates were highly similar; their RDFs, coordination numbers, and FFTs showed no evidence for crystallinity (Fig. 3c and 3d, Fig. S4f and S8a-d). Thus, despite the more solid-like characteristics of the P22L condensate at the mesoscale^16^, the Ape1 particles are locally amorphous, i.e. as in the liquid state.

To assess potential differences between WT and P22L condensate architecture in more detail, we extended our analysis to three-body correlations of Ape1 dodecamers. Specifically, we determined the distribution of the triplet angles between the vectors connecting an Ape1 dodecamer with two of its neighbours within the first coordination sphere (CS1) of the RDF (Fig. S8e). Unlike the maximally disordered Lennard-Jones (LJ) fluid, which exhibits a smooth angular distribution due to the lack of directional bonding or geometric constraints, our measurements revealed a different scenario for Ape1 condensates. There was a highly significant enrichment in triplet angles of ∼53°, while there was a clear depletion near ∼80° greater than 3 standard deviations of the LJ-fluid, which explains the tight packing needed for autophagic cargo assembly (Fig. 3e and Fig. S8f). Remarkably, the WT angle distribution was indistinguishable from the P22L, yet both sharply contrasted the triplet angles of 90° and 180° expected for a cubic crystal of tetrahedra involving two edge-to-edge pairs of propeptide interactions (Fig. S8g). These deviations indicate that the particle arrangement in our system is not governed by purely isotropic LJ-type interactions but is influenced by additional structural or interaction-specific constraints. Given the non-random orientations identified in our initial angular distribution, we performed STA using a larger box encompassing multiple Ape1 dodecamers. Although the reconstructions did not reach high resolution, likely due to a high degree of flexibility in the interaction between Ape1 dodecamers, the averages showed distinct trimer and tetramer species with 60 ± 8.9° (trimer) and 60 ± 4.8° (tetramer) triplet angles for the WT and 60 ± 7.4° (trimer) and 60 ± 4.7° (tetramer) for the P22L, in agreement with the angular distribution analysis (Fig. S2, S6 and S8h).

To assess connectivity between Ape1 dodecamers, we used the coordinates derived from STA to map the positions of the 12 propeptide exit sites on each dodecamer (Fig. 3f). We then analysed the pairwise distances between propeptides of neighbouring dodecamers within CS1, inferring physical connectivity by assuming that each propeptide interacts with its nearest neighbour. This geometric approximation allowed us to estimate local connectivity patterns across the condensate. The distribution of nearest-neighbour (d1), and second nearest-neighbour (d2) distances revealed a broad range, with d1 maxima at 3.46 ± 1.24 and 3.48 ± 1.33 nm and d2 maxima at 5.28 ± 1.09 nm and 5.34 ± 1.19 nm, for WT and P22L respectively (Fig. 3g).

Comparison with the estimated propeptide dimensions, based on conformational ensembles calculated from NMR chemical shifts, suggested that the compact organisation of propeptides arises both from overlap at the binding interfaces and from partial back-folding of the propeptides (Fig. S9d)^14^. Importantly, no significant differences in propeptide spacing or spatial arrangement were observed between the WT and P22L, indicating that both maintain similarly flexible geometrical arrangements (Fig. 3g). Additionally, the refined analysis revealed that, on average, each Ape1 dodecamer contacted about five neighbours in both WT (5.27 ± 1.73) and P22L (4.97 ± 1.81), rising to more than seven when including all molecules within CS1 (8.10 ± 3.26 for WT; 7.22 ± 3.21 for P22L) (Fig. 3h). Thus, P22L and WT condensates display similar local packing. As expected, dodecamers located at the condensate surface exhibited a much lower connectivity than those in the condensate core (Fig. 3i).

### Short transient α-helical structures in the propeptide tune condensate material properties

To explain the observed morphological changes in the P22L condensate at the mesoscale, we hypothesised that the P22L mutation alters the binding affinity between individual propeptides, leading to a change in viscosity compared to the WT condensate. To test this hypothesis, we examined propeptide self-interaction using AlphaFold predictions. The predicted coiled-coil interface between the helices of the WT propeptide extended over the first ∼20 residues (Fig. 4a), consistent with conformational ensembles determined based on previously acquired NMR chemical shifts, which showed that the first approximately 17 residues sample an α-helical conformation to an extent of nearly 100% at propeptide concentrations above 100 µM (Fig. S9a-c)^14^. The average Ape1 concentration within the condensates observed in situ was estimated at 327 ± 27 mg ml^-1^ for the WT and 309 ± 34 mg ml^-1^ for the P22L mutant, corresponding to > 450 µM in both cases, permitting N-terminal helix formation (Fig. S9e). AlphaFold predicted dimeric, trimeric, and tetrameric propeptide assemblies with high confidence, while pentamer and hexamer predictions showed low confidence scores (Fig. S9f and S9g). These predictions highlight the heterogeneity of propeptide-propeptide interaction stoichiometry facilitated by the hydrophobic interface contacts, likely contributing to the observed lack of long-range order within Ape1 condensates.

**Figure 4:**
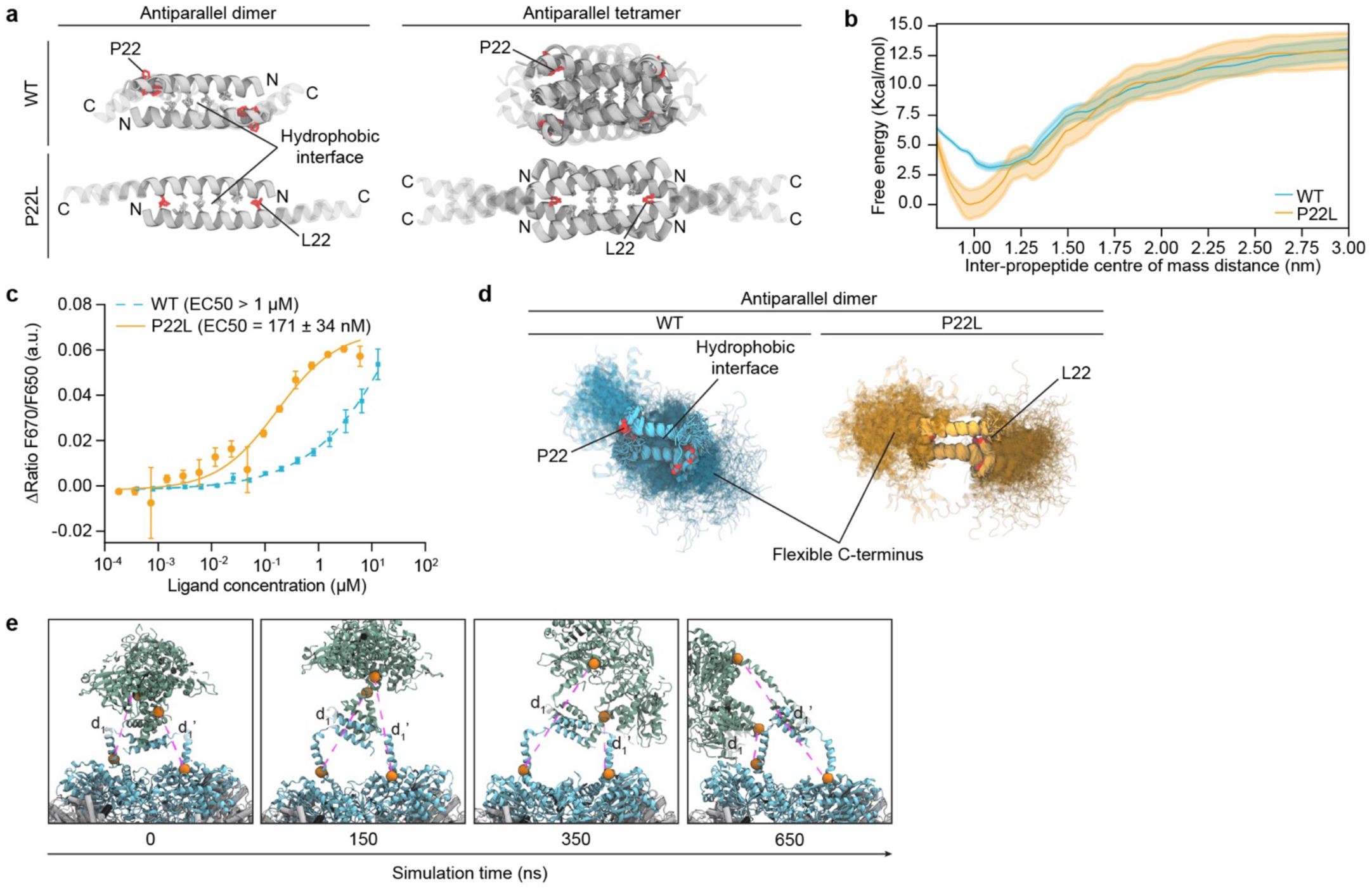
The P22L mutation increases propeptide-propeptide interaction strength. **a)** AlphaFold model renders of the best-scoring (highest interface pLDDT) antiparallel WT and P22L propeptide dimers (left) and tetramers (right) for increasing residue lengths (25, 30, 35, 40, and – for all except the P22L dimer – 45 residues). Residues 1-25 are shown as solid, and the remaining propeptide residues are transparent. Free energy profile as a function of the centre-of-mass distance between the N-terminal 30 residues of two propeptide chains for WT and P22L. **c)** Quantitative analysis of self-interaction of WT MBP-Ape1 (blue) and P22L MBP-Ape1 (orange). Spectral-shift dose response curves show higher self-interaction propensity of P22L as compared to WT. An EC50 value of 171 ± 34 nM for P22L is obtained, whereas WT did not reach saturation, but the value is likely to be higher than 1 µM. Data represent mean ± standard deviation calculated from at least 3 technical replicates. **d)** Superimposed snapshots from all-atom MD simulations of the antiparallel propeptide dimer for WT and P22L. **e)** Snapshots of all atom MD simulations for four full-length Ape1 chains in a tetrameric assembly taken at indicated timepoints.

Notably, the P22L mutation extended the helical interaction region in our AlphaFold predictions and increased the number of predicted interfacial contacts (Fig. 4a). The increased binding interface observed in the P22L mutant is consistent with enhanced binding affinity. This is supported by free energy profiles from umbrella sampling simulations, which revealed that the P22L dimer exhibits both a higher binding affinity and a larger energetic barrier to dissociation compared to the WT (Fig. 4b), indicating a more stable and long-lived interface. Furthermore, spectral shift analysis using purified recombinant WT and P22L Ape1 MBP-fusion proteins demonstrated a shift in binding propensity with EC50 values of 171 ± 34 nM for Ape1 P22L and above 1 µM for the WT (Fig. 4c, Fig. S9h and S9i).

To evaluate the stability and dynamics of the predicted interfaces, we performed all-atom molecular dynamics (MD) simulations of propeptide dimers and tetramers. In both WT and P22L variants, the coiled-coil interfaces remained stable, mediated by hydrophobic residues in the helical core, while flanking regions were highly dynamic, consistent with the variable distances observed in tomograms (Fig. 3g and 4d). MD simulations of a 2:2 Ape1 assembly further demonstrated the structural plasticity of these interfaces, with the propeptides sampling a broad range of conformations and distances (Fig. 4e, Fig. S9j and Supplementary Video S2).

Combined, these data suggest that the P22L mutation strengthens interparticle interactions by extending and stabilising the helical surface at the N-terminus, while the flexible C-terminus remains disordered, as in the WT. The increased binding affinity likely raises condensate viscosity, slowing internal diffusion, reducing exchange with the dilute phase, and delaying droplet shape recovery.

### Coarse-grained simulations establish a digital twin of condensate dynamics and assembly

To directly assess whether changes in propeptide interaction strength alone can account for the altered mesoscale morphology of P22L condensates observed in situ, we developed a coarse-grained simulation model of the Ape1 condensate. Each Ape1 dodecamer was represented as a sphere with 12 interaction sites located along the edges of a tetrahedral scaffold (Fig. 5a). Using LJ-like potentials for these patchy-particles, we tuned the site-site interaction strength to match experimental values of the density, relative binding affinity and number of neighbours for WT and P22L propeptides (Fig. S10a-d). Systematic screening of interaction parameters revealed a narrow window in which condensation is facilitated while particles remain distributed between the dense and dilute phase, allowing for particle exchange (Fig. S10d and S10e). Our model therefore predicts a small range of propeptide-propeptide binding affinities which support Ape1 condensate formation. Accordingly, sequence alignment analysis of Ape1 homologs belonging to diverse species of budding yeast shows conservation of the amino acids at the N-terminus of the propeptide, highlighting specific sequence requirements for transient structure formation of an interaction-mediating leucine-zipper-like structure (Fig. S10f). Contrastingly, the propeptide C-terminus displays large variation between species, suggesting that its disorder can be achieved by diverse sequence compositions. MD simulations of the WT system yielded spherical condensates with partial surface dynamics (Fig. 5b (top) and Supplementary Video S3). By contrast, simulations using stronger interactions for the P22L mutant produced irregular, non-spherical condensates with minimal particle exchange and a pronounced decrease of particles in the dilute phase, recapitulating the morphology changes observed in situ (Fig. 5b (bottom) and Supplementary Video S4).

**Figure 5:**
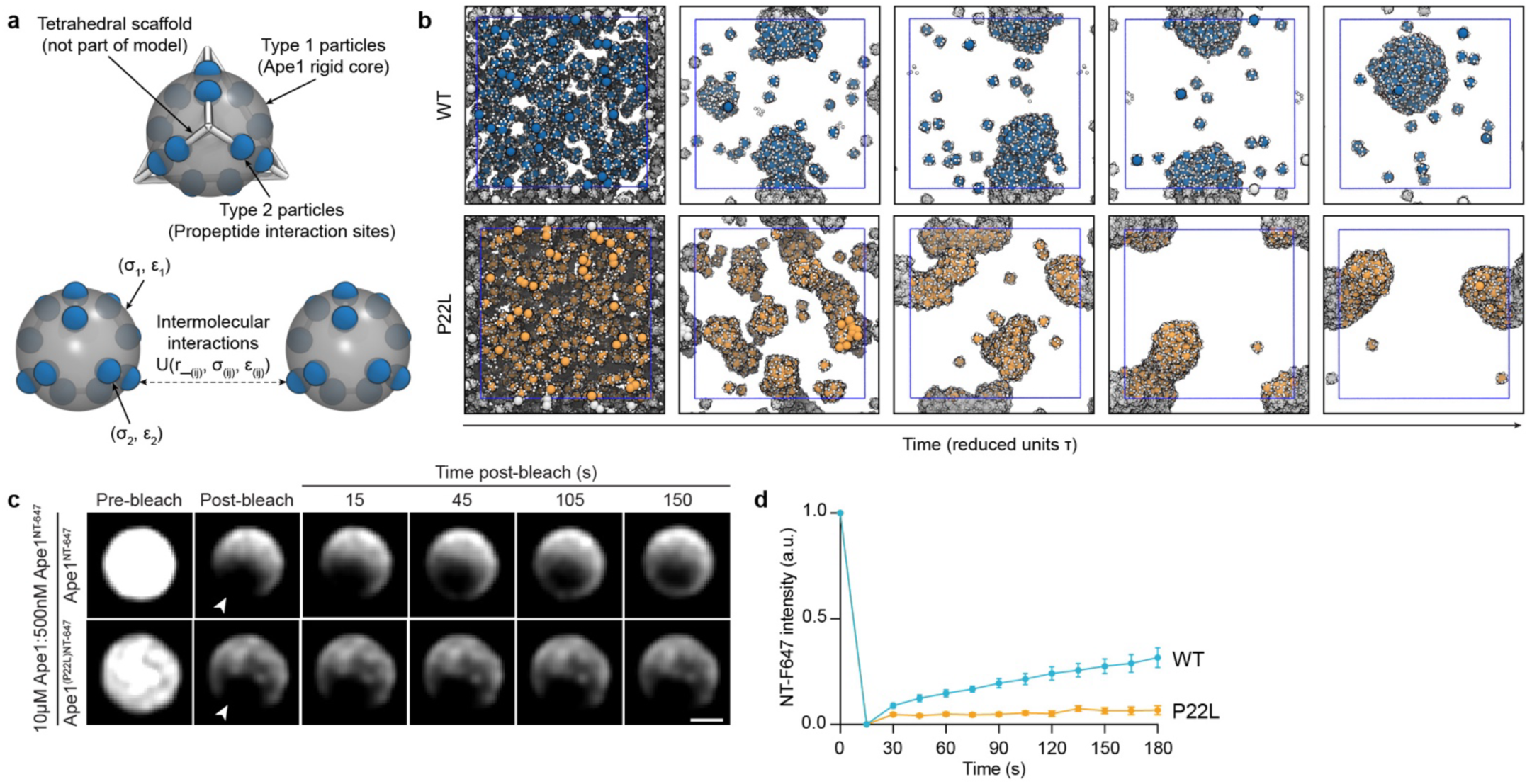
Propeptide binding affinity regulates Ape1 condensate dynamics. **a)** Schematic representation of coarse-grained simulation model with Ape1 dodecamers represented as spheres and 12 sticky patches located along the edges of a tetrahedral scaffold. Particles interact via the Lennard–Jones potential U, as defined in Methods eq. 6, with r_(ij) the distance between particles, and σ(ij), and ε(ij) the distance and energy scale for interactions between particle types i and j, respectively**. b)** Snapshots of coarse-grained simulation generated as in a) for interaction strengths approximated for WT (top) and P22L (bottom) propeptides. Representative deconvolved images of fluorescence recovery of Ape1 droplets containing WT unlabelled Ape1 with NT-647 dye-labelled WT (top) or P22L (bottom) Ape1 after photobleaching at the regions indicated by the white arrow. Scale bar = 1 µm. **d)** Quantification of fluorescence recovery at the surface of in vitro WT and P22L Ape1 droplets. Error bars represent the mean ± standard error of the mean for n = 10 structures per condition per replicate. 3 technical replicates were performed.

To test the prediction that particle exchange is reduced in the P22L mutant, we performed fluorescence microscopy on droplets formed with recombinantly purified Ape1 (Fig. S10g and S10h). While WT Ape1 readily formed spherical droplets in vitro, P22L Ape1 formed irregularly shaped condensates, consistent with previous observations using RFP-labelled proteins (Fig. S10i)^16^. To directly test whether the P22L mutation reduces particle exchange between condensates and the dilute phase, we reconstituted Ape1 droplets by combining unlabelled WT Ape1 with a 20-fold dilution of fluorescently (NT-647) labelled Ape1, either WT or P22L (Fig. S10j). This design ensured comparable droplet formation in both conditions while allowing us to track the behaviour of WT and P22L Ape1 particles. Fluorescence recovery after photobleaching (FRAP) experiments revealed 3-fold higher recovery of outer versus interior WT molecules of Ape1 droplets, consistent with higher Ape1 exchange with the dilute phase at the condensate edge (Fig. 5c and 5d, Fig. S10k and S10m). By contrast, we did not detect recovery of P22L at the edge or interior of Ape1 droplets during the FRAP experiment (Fig. 5c and 5d, Fig. S10l and S10m), indicating that the P22L Ape1 at the surface of droplets does not exchange with the dilute phase within the timeframe of the experiment.

Together, these results demonstrate that an increase in propeptide binding affinity, driven by a single amino acid substitution, is sufficient to shift condensate behaviour towards a highly viscous, glass-like state. This explains not only the observed loss of material fluidity due to an increase in the site-specific propeptide-propeptide interface stability but also the morphological deviations of the P22L mutant from the spherical arrangement (Fig. 6).

**Figure 6:**
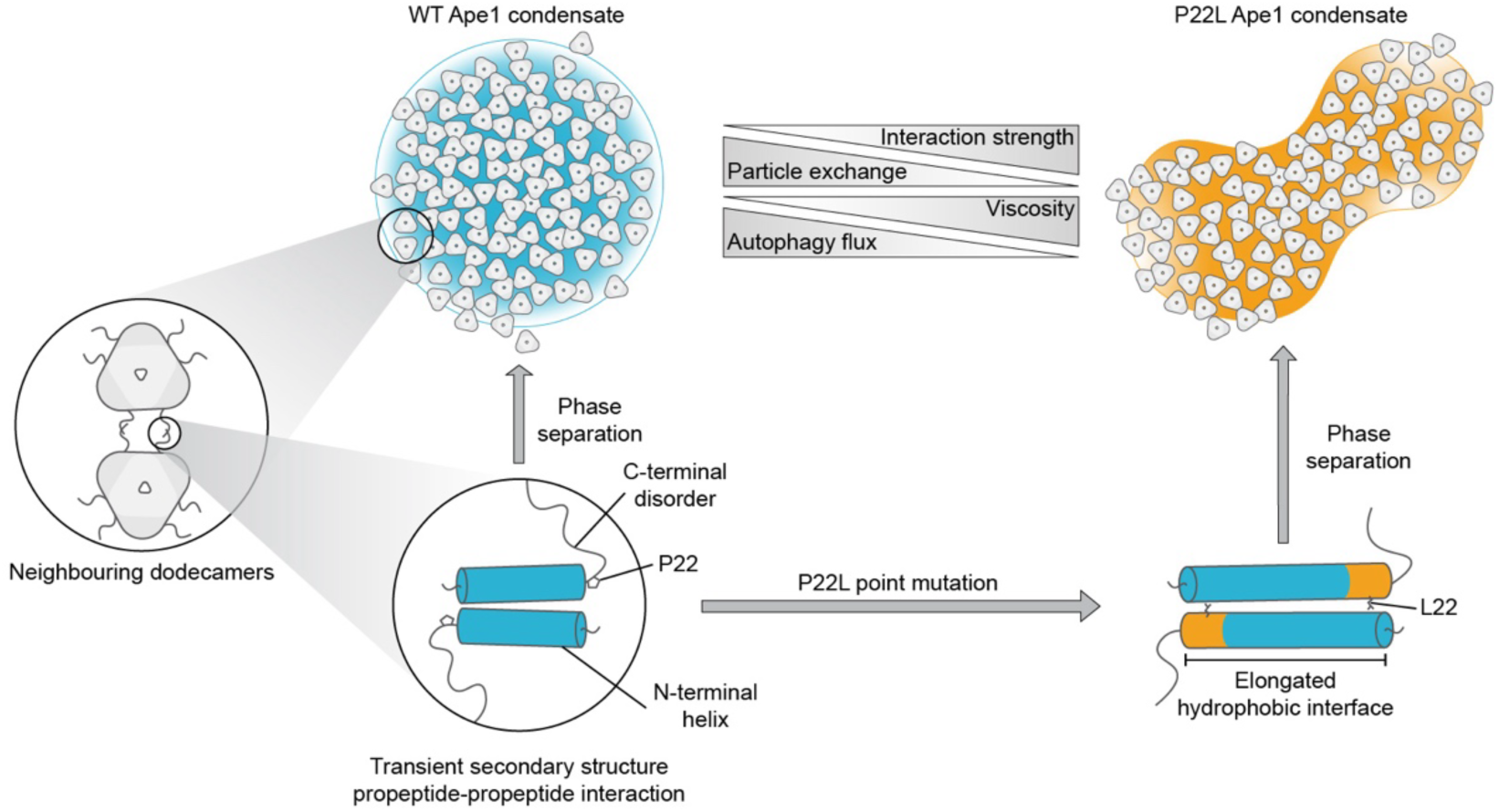
Model for mechanism of phase separation in Ape1 condensates. Ape1 condensate formation is driven by hydrophobic interactions between transient α-helices at the propeptide N-terminus. Upon introduction of the P22L mutation, an extension of the hydrophobic interface increases the propeptide binding affinity, causing changes to condensate dynamics and morphology.

## Discussion

Although condensate material state has long been linked to cellular function, our work establishes at molecular resolution how a single amino acid substitution can strengthen intermolecular interactions between intrinsically disordered regions (IDRs), drive a liquid-to-glass transition, and, in turn, abolish autophagic clearance. This establishes a direct link between molecular interaction strength, condensate material properties, and physiological function.

Our in situ architectural model of the Ape1 condensate at high resolution, spanning from mesoscale droplet organisation to the molecular determinants of condensate material properties (Fig. 6), reveals how these changes can be understood within the sticker-spacer framework of associative polymers^1,37,38^. Specifically, the N-terminal 17 residues, which adopt a short alpha-helical conformation, function as the "sticker," while the adjacent low-complexity region, enriched in charged residues (glutamate and lysine), serves as the "spacer." Consistent with this model, the P22L mutation, which increases the hydrophobic binding interface between helical stickers, enhances binding affinity, whereas mutations that disrupt the hydrophobic interface, such as L11S, impair both condensate formation and autophagic delivery^13,14,34,36^. This discovery highlights how transient secondary structure elements can act as stickers, with their interaction affinity serving as a critical determinant of condensation. In line with our observations, a transient short α-helical element in the CTD of the disease-relevant protein TDP-43 plays a major role in its phase separation behaviour. Consistently, helix-stabilizing mutations further enhance phase separation in vitro^39,40^. Beyond classical physicochemical parameters (e.g., charge, dipole moment, π–π stacking), our findings highlight a concentration-dependent mechanism of interaction, wherein transient secondary structures enable site-specific, low-affinity contacts to tune condensate behaviour. This mechanism may account for the specificity in phase-separating protein assemblies, broadening the molecular grammar that governs phase separation^41^.

Recent artificial protein design using RoseTTAFold^42^ identified well-folded proteins that interact with specific conformations of otherwise IDRs in phase-separating proteins, thereby disrupting condensate formation^43^. Such an induced-fit mechanism, where the binder selects a single conformation from a broad ensemble, seems to be common in interactions between folded proteins and IDRs^44,45^. Multivalency is further amplified by oligomerization of Ape1 monomers into dodecamers, effectively forming an emergent sticker^1^. Such emergent stickers, arising through clustering or microphase separation, represent a general mechanism for enhancing phase separation through increased valency, as supported by impaired Ape1 condensation upon disruption of dodecamer assembly^30^.

Importantly, enhancing sticker interaction strength upon P22L mutation alters the condensate’s material state without perturbing the relative organisation of Ape1 dodecamers. The resulting assemblies exhibit characteristics of colloidal glasses, amorphous in structure but markedly viscous, with arrested dynamics, suggesting a kinetically trapped system due to elevated binding affinities. These findings propose a broader model for condensate regulation, highlighting how molecular interaction strength influences glass-like dynamics in biomolecular assemblies. This tuneable viscosity may underlie functional adaptability in phase-separated compartments.

Functional Ape1 condensates are recognized by selective autophagy receptors, engulfed in autophagosomes, and delivered to the vacuole. Cargo selectivity depends on forming a tight meshwork that excludes non-cargo proteins. Our analysis revealed that Ape1 condensates are viscous, densely packed assemblies with minimal space between dodecamers, explaining cytosolic protein exclusion. Cryo-ET and fluorescence imaging further demonstrated that condensation excludes small proteins, reinforcing its role in defining discrete, selective cargo.

Although receptor recruitment occurred in both WT and P22L condensates, the rigid P22L surface was associated with impaired subsequent phagophore engagement, highlighting that receptor mobility and surface fluidity, rather than receptor binding per se, govern productive autophagic trafficking^16,25^. This aligns with observations that amorphous aggregates, which maintain fluid-like surfaces, are readily degraded, whereas rigid, low-mobility amyloid fibrils resist autophagic clearance^46^.

Together, these findings underscore how the physical state of condensates governs their biological fate and highlight the need to interrogate condensate material properties as key determinants of both function and dysfunction. Condensate material state emerges as a decisive control point, where even a single amino acid can tip the balance between dynamic function and pathological arrest.

## Materials and Methods

### Yeast growth conditions

Yeast strains used in this study are listed in Supplementary Table S1. Dense stationary phase yeast cultures supplemented with glycerol for cryo-protection were stored at -80° and streaked onto fresh agar plates with required auxotrophic selection prior to inoculation of liquid cultures for experiments. Fluorescent tagging and deletion of genes at endogenous loci for strains constructed as part of this study was performed by PCR-based genomic integration^48^. Generation of centromeric yeast expression plasmids listed in Supplementary Table S2 was performed using standard cloning and site-directed mutagenesis techniques. Yeast strain genotypes were confirmed by genomic PCR, immunoblotting or live cell microscopy. Unless otherwise stated,

*S. cerevisiae* cells were cultured in synthetic medium (SD, 0.17% yeast nitrogen base without amino acids and ammonium sulphate, 2% glucose and amino acids as required for selection of transformed centromeric plasmids) at 30°C shaking at 220 rpm. For induction of Ape1 overexpression in strains containing Ape1-encoding plasmids, a final concentration of 250 µM CuSO4 was added to cultures at OD600 ≤ 0.1 and cultures were grown to mid-log phase. Autophagy induction was achieved by treatment of mid-log phase cultures with a final concentration of 100 nM rapamycin for 3 hours.

### Live cell fluorescence microscopy

#### Sample preparation and data acquisition

For live cell fluorescence microscopy, yeast cultures were grown in low fluorescence synthetic medium (yeast nitrogen base without amino acids, folic acid and riboflavin, 2% glucose and amino acids as required for selection of transformed centromeric plasmids). Cultures were grown to mid-log phase with the exception of the mScarlet exclusion experiment (Fig. 2e and 2f), for which cells were grown to OD600 > 8.0 to induce formation of larger Ape1 condensates, as performed previously^16^. Where required, cells were treated with CuSO4 and rapamycin as described above to induce Ape1 overexpression and autophagy then applied to microscopy slides pre-treated with 1 mg ml^-1^ concanavalin A type IV. Imaging was carried out on a Nikon Eclipse Ti2 inverted microscope operated by NIS-Elements software version 5.10 (Nikon Instruments) with an Apo TIRF 100x/1.49NA oil objective and a Hamamatsu C11440-22C CMOS camera. Images were deconvolved using the theoretical point spread function (PSF) and Classic Maximum Likelihood Estimation (CMLE) algorithm in Huygens Professional (version 24.04, Scientific Volume Imaging, The Netherlands, http://svi.nl/).

#### Image analysis

For automated quantification of BFP-Ape1 and sfGFP-Atg19 puncta abundance in WT and P22L Ape1 overexpression strains, yeast cells in central slice brightfield images were detected using YeastMate (https://github.com/hoerlteam/YeastMate/). Fluorescent puncta were detected by threshold-based segmentation in maximum intensity projections of deconvolved BFP or GFP channel images in Fiji^49^ and puncta per cell were counted using a custom Python script. For quantification of mScarlet exclusion from Ape1 condensate interiors in deconvolved microscopy images, line profiles were drawn through the central slice of all visible sfGFP-Atg19 rings and fluorescence intensity of both sfGFP-Atg19 and mScarlet signal were measured using the “Plot Profile” tool in Fiji^49^ then normalised to the maximum intensity value in each channel. Maxima of sfGFP-Atg19 signal were detected and used to define “inside” and “outside” regions for calculating mean mScarlet fluorescent intensity along the line profiles.

### Immunoblotting

Yeast cell lysates from 1.0 OD600 were prepared via trichloroacetic acid (TCA) precipitation. Cells were harvested at 2000g for 5 min and cell pellets were snap frozen in liquid nitrogen then resuspended in 1 ml ddH2O and 150 µl of 1.85 M NaOH with 7.5% β-mercaptoethanol was added for 15 min on ice. 150 µl 55% TCA was added for 10 min on ice and precipitated proteins were pelleted at 21130g for 2 min at 4°C and all remaining TCA was removed. Protein pellets were resuspended in HU buffer (8 M urea, 5% SDS, 200 mM Tris-HCl pH 6.8, 1 mM EDTA, 100 mM DTT, bromophenol blue) and incubated at 65°C and 1000 rpm for 10 min. Proteins were separated by NuPAGE^TM^ 4-12% Bis-Tris gels (Invitrogen) and transferred to low fluorescence PVDF membranes (Immobilion®-FL). Imaging of membranes and analysis of signal from fluorescent secondary antibodies was performed using an Odyssey® Fc imager operated by Image Studio (version 5.2, LI-COR).

### Antibodies

The following antibodies were used in this study: rabbit polyclonal anti-Ape1 (1:3000 dilution, provided by Dr Thomas Wollert), mouse monoclonal anti-Pgk1 (1:10000 dilution, clone 22C5D8, ref no. 459250, lot no. K4823, Novex Life Technologies), IRDye 680RD Goat anti-rabbit (1:15000, ref no. 926-68071, lot no. D10603-15, Licor), IRDye 800CW Goat anti-mouse (1:15000, ref no. 926-32210, lot no. D10825-15, Licor).

### Cryo-electron tomography (cryo-ET)

#### Grid preparation

Yeast cells were grown as described above with induction of Ape1 overexpression using 250µM CuSO4 and 3 hours of 100nM rapamycin treatment. Cultures were then diluted to an OD600 of 0.8 for use in the cryo-ET workflow. For correlation of giant Ape1 condensates in sfGFP-Atg19 strains, autofluorescent 1 µm diameter Dynabeads (Dynabeads MyOne carboxylic acid No. 65011, Thermo Fisher Scientific) were added to cultures at a 1:20 dilution. Prior to vitrification, EM grids (200 Mesh Au SiO2 R1/4, Quantifoil) were glow discharged for 90 s on both sides using a Pelco easiGlow device. Vitrification was carried out using an EM GP2 grid plunger (Leica Microsystems) with chamber temperature of 20°C and 90% chamber humidity. 3.5 µl of cell suspension was applied to grids and back blotting was carried out for 3 s followed by plunge freezing in liquid ethane at -184°C. Grids were clipped into autogrid rings modified for cryo-FIB milling and stored in liquid nitrogen until further use.

#### Cryo-confocal fluorescence microscopy

Grids of sfGFP-Atg19-positive cells were imaged using a TCS SP8 Cryo-CLEM (Leica Microsystems) with a 50x/0.9NA air objective operated by LAS X Navigator software (version 3.5.7, Leica Microsystems). Grids were mapped by collecting image stacks at 2x2 binning with 1.5 µm z-spacing in brightfield and widefield fluorescence modes. Grid maps were used to identify target grid squares. Confocal image stacks of target grid squares were acquired at 488 nm laser excitation with z-spacing of 300 nm and xy pixel spacing of 84.4 nm. Cryo-confocal fluorescence image stacks were deconvolved using the theoretical PSF and CMLE algorithm in Huygens Professional (version 24.04, Scientific Volume Imaging, The Netherlands, http://svi.nl/) and resliced in Fiji^49^ to achieve cubic voxels.

#### Focused ion beam (FIB)-milling

Focused ion beam (FIB)-milling was performed using an Aquilos2 dual-beam cryo-focused ion beam microscope (cryo-FIB) (Thermo Scientific) equipped with cryo stage cooled to < -180°C. Grids were coated using the organoplatinum gas injection system for 30-60 s followed by sputter coating with elemental platinum for 20 s. Mapping of grids by scanning electron microscopy (SEM) and correlation with previously acquired widefield fluorescence grid maps was carried out using MAPS software (version 3.29, Thermo Scientific). Lamella sites were placed at positions where cryo-confocal fluorescence stacks had been acquired. Milling was performed at a milling angle of 8°. To mill lamella at the z-height of the target sfGFP-Atg19 fluorescence signal, 3D correlation between cryo-confocal fluorescence data and FIB images of the target cell clump taken at the milling angle was carried out using 3D-Correlation Toolbox (https://3dct.semper.space/)^50^ using Dynabeads visible in both the fluorescence and FIB imaging modalities as fiducials. For this, approximately 8 fiducials were selected to calculate the target signal location for each lamella. Milling patterns were placed manually according to the correlation output to capture the target fluorescent signal within the lamella. Automated FIB-milling was carried out using AutoTEM Cryo software (version 2.4, Thermos Scientific) with a step-wise decrease of milling pattern distance and ion beam current to achieve a target lamella thickness of ∼150 nm. Following automated milling, manual polishing was carried out to further reduce lamella thickness. Lamella quality and thickness was evaluated by SEM imaging of the final lamellae at 3kV.

#### Tomography data acquisition

Tilt series acquisition was carried out on a Titan Krios G4 (Thermo Scientific) cryo-transmission electron microscope operated at 300 kV with Selectris X energy filter and Falcon 4i direct electron detector (Thermo Scientific). Firstly, montaged grid maps and lamella maps were acquired at 125x and 8700x magnification respectively. Additionally, images of grid squares with milled lamella were acquired at 470x magnification. For tilt series acquisition site selection, 2D correlation was carried out between the TEM grid square images and fluorescence stacks using 3D-Correlation Toolbox, using visible Dynabeads and prominent lamella features as fiducial markers. Tilt series images were collected using SerialEM (versions 4.0-4.2)^51^ at 64000x magnification, corresponding to a calibrated pixel size of 1.895 Å. The energy slit was set to 10 eV and a dose-symmetric acquisition scheme was used with tilt range of -60 to +60°, tilt increment of 2°, target defocus range of -1.5 to -4.5 µm and total dose of ∼150 e^-^/Å^2^. 10 frames per tilt were acquired and aligned on-the-fly in SerialEM. Data acquisition parameters are also detailed in Supplementary Table S3.

#### Tomogram reconstruction and segmentation

Acquired tilt series were filtered by cumulative electron dose in MATLAB as previously described^52^ and per-tilt contrast transfer function (CTF) estimation was performed using gctf (version 1.06)^53^. Bad tilts were discarded upon visual inspection and tilt series alignment by patch tracking and weighted back projection reconstruction at bin4 using 10 iterations of SIRT-like filtering was carried out using AreTomo2 (version 1.0.0)^54^ and IMOD (version 4.12.62)^55^, respectively. Segmentation of cellular membranes was performed on fSIRT tomograms using MemBrain-Seg (https://github.com/teamtomo/membrain-seg/)^56^ , manually refined in Amira (version 2023.2, Thermo Scientific) and displayed using the ArtiaX plug-in (https://github.com/FrangakisLab/ArtiaX/)^57^ in ChimeraX (version 1.9)^58^. Denoising was performed on selected tomograms for display in figures. For this, frames were split into even/odd batches and aligned using IMOD. Tomograms were reconstructed at bin4 via weighted back projection for odd and even frames using alignment parameters previously calculated by AreTomo during full tomogram reconstruction. Odd and even tomograms were used to train a Cryo-CARE (https://github.com/juglab/cryoCARE_T2T/)^59^ model for denoising which was applied to the full reconstruction of each tilt series.

### Subtomogram averaging

Template matching was used to identify the exact positions and orientations of Ape1 dodecamers within Ape1 condensates. First, 4 Ape1 WT CTF-corrected tomograms were reconstructed at bin8 with Warp^60^ using the alignment parameters from the initial AreTomo reconstructions. Template matching of these tomograms was initially performed using Dynamo^61^ with a cone range of 360 degrees with cone sampling of 5 degrees and in plane range of 360 degrees with in plane rotation of 10 degrees. The cryo-EM map (EMD-30652) of *Schizzosaccharomyces pombe* Ape4^28^ was rescaled using EMAN2^62^ and used as the template together with a spherical mask created using Dynamo^61^. From the template matching results, 12000 subtomograms were extracted at bin4 using Warp and classified using RELION (version 3.1.3)^63^. A class of 2475 particles resembling Ape1 was selected and corresponding subtomograms were re-extracted at bin2, classified and refined into a 25 Å average.

Using the initial 25 Å Ape1 dodecamer average as a template, high confidence template matching was subsequently performed using GAPSTOP™^64^ based on the STOPGAP^65^ software package similarly to previous studies with the following parameters: 5° angular step, low-pass filter radius = 15, high-pass filter radius = 1, apply_laplacian = 0, noise_correlation = 1 and calc_ctf = 1. The above optimized settings produced distinguishable peaks when applied to bin2 tomograms as they were visualised in napari (https://github.com/napari/napari/)^66^ (Fig. S2). To ensure all particles were extracted, we visually inspected each tomogram and adjusted the z-score value between 15-17 units for the extraction of the coordinates. This selection was based on the constrained cross-correlation (CCC) value from template matching and each tomogram was visually inspected for correct coordinate extraction.

These coordinates were subsequently extracted as subtomograms in Warp^60^. 3D classifications (classes = 4, T = 0.5, iterations = 35, mask set to 220 Å) were performed in RELION (version 3.1.3)^63^ to clean particle lists, followed by 3D refinement to curate particle poses. Ape1 WT was initially refined in C1 symmetry and then refined to 3.84 Å) using T symmetry. For the dataset of Ape1 P22L in the presence of endogenous WT Ape1, subtomograms were initially extracted at bin2 and then selected subtomograms were re-extracted and refined at bin1 to a final global resolution of 5.6 Å. For the dataset of Ape1 P22L in the absence of endogenous WT Ape1, subtomograms were extracted from two independent runs of template matching over different tomograms and subjected to 3D classification in RELION. Coordinates corresponding to intact Ape1 dodecamers were then selected and merged at bin2, and refined using T symmetry to a global resolution of 8.8 Å. Further cleaning using 3D classification and re-extraction of these subtomograms at bin1, did not improve the resolution, with the best reconstruction reaching a resolution of 9.1 Å. For all final maps in this study, post-processing was performed using a calibrated pixel size of 1.895 Å/px. Further details of the subtomogram averaging pipeline for Ape1 dodecamers are shown in Supplementary Table S4.

To observe Ape1 particles with their neighbours as subtomogram averages, we extracted bin2 subtomograms at a larger box size (120 pixels) (Fig. S2 and S6). Classification in RELION revealed three main types of Ape1 arrangements: two, three and four Ape1 particles. Comparison of the local Ape1 particle distribution within these classes with the global arrangement across the whole condensate was performed by determining coordinates of the centre of mass for each density and determining the Euclidean distance between these coordinates, as well as the inter-vector angle between all combinations of 3 neighbouring Ape1 particles within the classes.

### Ape1 atomic model building and refinement

We observed a good fitting of the previous crystal structure of prApe1 L11S (PDB 5JH9)^14^ into our electron density map based on rigid body fitting performed in ChimeraX. Nevertheless, a complete model of the dodecamer was built from AlphaFold2^67^ predictions of the prApe1 monomer including the propeptide. The model was then refined into the map by using ChimeraX’s function *fit in map* and ISOLDE (version 1.9)^68^. After running ISOLDE, the model was inspected in Coot (version 0.9.8.95 EL)^69^ where non-fitting residues were removed. Then, a second refinement was performed using Phenix^70^. ChimeraX (version 1.9) was used to visualise EM maps and models^58^.

### Ape1 condensate sphericity analysis

Circularity analysis was performed on all bin4 fSIRT reconstructed tomograms in which the Ape1 condensate did not exceed the tomogram field of view in the xy plane. Ape1 condensates in three slices per tomogram (the central slice and slices at positions +20 and -20 voxels from the central slice) were manually segmented in Dragonfly ORS (version 2024.1, http://www.theobjects.com/dragonfly, Comet Technologies Canada Inc.). Circularity of Ape1 condensates was measured from these segmentations by 3 orthogonal methods. Firstly, circularity (C) was calculated directly from the segmentation area (A) and perimeter (p) using the formula:

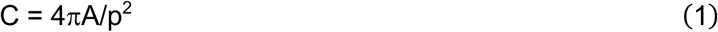

Secondly, ellipse fitting of segmentations was performed using the Python OpenCV package (version 4.11.0) and the minor/major axis ratio was calculated where a ratio of 1.0 is equivalent to a perfect circle. Thirdly, correlation between the mean curvature of the segmentations and the curvature of an area-equivalent circle was calculated. To calculate mean curvature of the of the segmentations, contours were extracted and approximated, allowing for an error of up to 2% of the total perimeter. Curvature at each contour point was calculated using the Menger curvature formula:

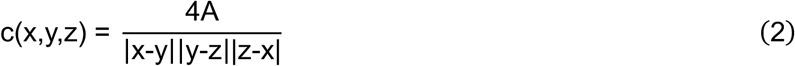

where x, y and z are consecutive contour points, and A is the area of the triangle enclosed by these points. Curvature of circles with area equal to the segmentations was taken to be 1/r, where r is the radius of the circle.

### Curation of Ape1 particle lists

To analyse the spatial organisation and potential interactions of Ape1 particles, we used the positions and orientations derived from subtomogram averaging performed with RELION (version 3.1.3)^63^. These coordinates served as the basis for subsequent spatial analyses across the datasets. During subtomogram averaging, only particles with high resolution features were preserved. However, 3D classification results showed a larger number of particles with features consistent with the Ape1 dodecamer structure. To generate more complete Ape1 particle lists, we selected all the classes that resembled the Ape1 dodecamer and removed duplicate particles using a 110 Å threshold. Particles with two or more neighbours within a distance of the first peak of the RDF (15 nm) were then selected as part of the Ape1 condensates. Finally, manual inspection of these particle lists using the ChimeraX (version 1.9) ArtiaX plug-in^57,58^ allowed us to clean the lists further for remaining particles that did not form part of the Ape1 condensate. For the particles from tomograms of the Ape1 P22L strain with no endogenous WT Ape1, coordinates at bin2 were used for analysis since the resolution did not reach Nyquist value at bin2 and the particle poses could not be further refined by extracting and refining the particles at bin1.

### Radial and angular distribution functions

To assess the spatial organisation of Ape1 particles and analyse their local structural arrangement, we first computed the radial distribution function (RDF). We report the normalised RDF, g(r), defined as the ratio:

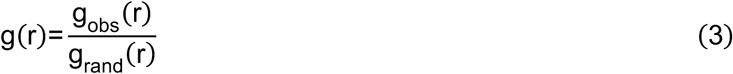

where g_obs_(r) is the observed distribution of pairwise distances between Ape1 particles (obtained from a histogram of inter-particle distances), and g_rand_(r) is the expected distribution derived from randomly placed particles within the same volume, generated using Trimesh (Dawson-Haggerty et al., 2019; https://trimesh.org/). The reference volume was defined by the concave hull enclosing the observed

Ape1 particle cloud. The Ape1 condensate boundary was approximated by a 3D Delaunay triangulation of particle coordinates using PyVista^71^. The alpha parameter was tuned to the first minimum of the surface-to-volume ratio of the resulting mesh. Hulls were exported using Trimesh and visualised together with particle coordinates and averages via Molecular Nodes (version 4.4.3)^72^ in Blender (Blender Foundation, 2025; https://www.blender.org/).

We then analysed the local angular arrangement of neighbouring Ape1 particles by computing the angles between vectors connecting each particle to its nearest neighbours. For each central particle, neighbours were identified within the CS1 (Fig. S8e). Next, we calculated vectors v_i_ from the central particle to each of its neighbours i. For every pair i,j of these neighbouring vectors, we computed the cosine of the triplet angle θ_ij_ as:

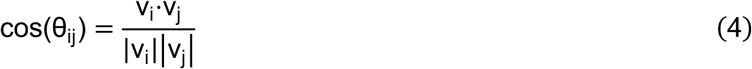

This analysis produced distributions of cosines of triplet angles around a central particle (Fig. 3e), enabling us to quantify local spatial ordering and detect potential patterns among adjacent particles. Note that for particles randomly placed on a sphere around the central particle, cos(θ) will be distributed uniformly with a sharp onset at small angles due to steric exclusion. To obtain a truly random reference for the angular arrangement of particles, we generated coordinates from an all-atom MD simulation of Argon atoms in the NVT ensemble, as described below. To quantify deviations between observed and simulated angle distributions, we calculated standardized residuals. Specifically, the mean angle of the observed dataset (μ_obs_) was compared to the mean of the simulated distribution (μ_sim_), and the difference was normalized by the standard deviation of the simulation (σ_sim_):

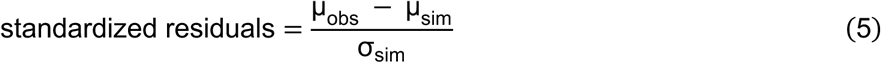

### Local concentration estimation

Local concentration was estimated by identifying the 32 nearest neighbours of each Ape1 particle, corresponding to the rounded second coordination shell. The local volume was defined as a sphere centred on the particle with a radius equal to the distance to its 32nd neighbour, and the number density was calculated as the neighbour count divided by this volume. Mass density was obtained from the molecular weight of the Ape1 dodecamer (12 × 57,093 Da). The resulting distributions were plotted as histograms and fitted with a two-component Gaussian mixture model (GMM), representing a high-density “internal” fraction and a lower-density “surface” fraction. To minimize artifacts from particle picking, only the best-picked tomograms were analysed from the WT and P22L datasets, as determined by clear visual separation of both Gaussian peaks.

### Propeptide connectivity analysis

To assess potential connectivity between the propeptides of neighbouring Ape1 particles, we analysed the spatial arrangement of residue 40 from each chain, which marks the start of the propeptide segment.

#### Mapping propeptide positions

For each Ape1 particle, we calculated the 3D vector displacement from its centre to residue 40 in each of its 12 chains. Using these displacements, we mapped the position of each particle (as obtained from subtomogram analysis) to the coordinates of its 12 corresponding propeptide start sites. This yielded a new dataset of propeptide positions in the cellular tomograms.

#### Identifying neighbouring propeptides

Using cryoCAT (https://github.com/turonova/cryoCAT/), we identified all neighbouring Ape1 particles within a 21.2 nm (the first minimum of the RDF) radius of a given central Ape1 particle. For each of these neighbours, we retrieved the positions of their 12 propeptide sites, resulting in a local environment of propeptides surrounding the central particle.

#### Determining connectivity

For each of the 12 propeptides of the central Ape1 particle, we computed the distances to all propeptides of neighbouring particles. Each propeptide was assumed to connect to its nearest neighbour. Two Ape1 particles were considered “connected” if they shared at least one pair of propeptides linked in this way.

This method enabled us to infer pairwise connectivity between particles based on their local propeptide geometry in situ.

#### AlphaFold modelling

We performed a systematic AlphaFold 2.3 Multimer^67,73^ screen of different oligomeric assemblies of Ape1 propeptide constructs. In detail, we ran predictions of 5 different oligomerization states (dimers, trimers, tetramers, pentamers, and hexamers) with 5 different propeptide sequence lengths (residues 1-25, residues 1-30, residues 1-35, residues 1-40, residues 1-45) for WT and P22L Ape1, respectively (50 different predictions in total). We used a local installation of AlphaFold 2.3 Multimer with AlphaPulldown 0.30.7^74^ to run the predictions with separated multiple sequence alignment (MSA) and prediction steps. Since we ran predictions of varying homooligomers of the same sequence we could reuse the single chain MSAs of each length for all 5 oligomerization states. For each system, we obtained 25 predicted structures (5 predictions per AlphaFold 2.3 model). We ran with 10 recycles and did not relax predicted structures.

We analysed the interfaces of the sequence part shared by all constructs: the first 25 residues. We computed the interface pLDDT score^75,76^ as the average residue pLDDT of all residues (in the range of 1 to 25) with a heavy atom within 5 Å of a heavy atom from a different chain (again, only considering residues 1 to 25). Furthermore, we sorted all predicted structures depending on the relative orientation of the first 25 residues between the different chains. Since all predictions consistently predicted a majority of these residues to be α-helical, we distinguished antiparallel and parallel arrangements between different chains. For any two chains termed A and B, we computed two Cα distances: (i) between residue 1 of chain A and residue 1 of chain B, and (ii) between residue 1 of chain A and residue 25 of chain B. If for any combination of chains distance (ii) was shorter than distance (i), we considered the arrangement to be antiparallel and otherwise to be parallel.

We then utilized AlphaFold 3^77^ to model the tetramer propeptide interface in the context of the full-length Ape1 chain. As input, we used two full-length Ape1 chains and two Ape1 propeptide (residues 1-25) chains. For the WT, we obtained 5 models from 1 seed using the AlphaFold 3 webserver. For the P22L, we obtained 50 models from 10 seeds using the webserver, and an additional 500 models from 100 seeds using a local AlphaFold 3 installation. For both systems, we selected a high-scoring model with the same interface as in the AlphaFold 2.3 Multimer predictions.

### All-atom molecular dynamics simulations

We used all-atom MD simulations to assess the structural stability of AlphaFold-predicted propeptide models, to compute the free energy profile between two interacting propeptides as a function of their separation, and to evaluate the relative flexibility of assemblies comprising two and four connected propeptides. Additionally, we used the same framework to analyse the triplet angles between randomly distributed particles in maximally disordered LJ-fluid, for comparison with our experimentally determined angular distribution of Ape1 particles.

#### Structural stability of the propeptides

All simulations were performed using GROMACS (version 2024.3)^78^. Proteins were modelled with the a99SB-disp force field, and water was represented by the matching a99SB-disp water model^79^. A summary of the simulated systems is shown in Supplementary Table S5. Each system was solvated in a triclinic dodecahedral box, ensuring a minimum of 1.2 nm between the solute and the box boundary. Periodic boundary conditions were applied in all directions. Long-range electrostatics were treated using the particle-mesh Ewald (PME) method^80^ with tin-foil boundary conditions, a Fourier grid spacing of 0.12 nm, and fourth-order interpolation. Coulomb and Lennard-Jones interactions were truncated at 1.2 nm. Bond constraints involving hydrogen atoms were enforced using the LINCS algorithm^81^, allowing a 2 fs integration time step.

Before the production runs, all systems were subjected to the following energy minimization and equilibration protocol. Energy minimization was performed using the steepest descent algorithm for up to 50 000 steps. Later, the system equilibration was conducted in two phases. First, a 1.0 ns simulation in the NVT ensemble was performed (constant volume and temperature = 300 K). This was followed by an additional 1.0 ns simulation in the NPT ensemble (constant pressure = 1 bar, and temperature = 300 K). During both equilibration phases, positional restraints (1000 kJ mol⁻¹ nm⁻^2^) were applied to heavy atoms of the protein. Temperature was controlled using the velocity rescaling thermostat with a stochastic term^82^, and pressure was maintained using the Parrinello–Rahman barostat^83^ with coupling constants of 0.1 ps (temperature) and 5.0 ps (pressure). After equilibration, position restraints were removed, and production simulations proceeded in the NPT ensemble using the same thermostat and barostat settings as above.

To assess the stability, flexibility, and organisation of the predicted Ape1 structures, we performed unrestrained all-atom MD simulations for both the WT and P22L Ape1 propeptides. For each variant, we simulated full-length propeptide dimers starting from the AlphaFold-predicted models (3 replicates of 1 μs each; total 6 μs). In addition, we simulated tetrameric assemblies containing four complete chains, again using three 1 μs replicates per variant. For the dimers and tetramers, the RMSD of the interface residues (residues 1 to 25 of each chain) reached a plateau after a few nanoseconds (10-50 ns), with small fluctuations thereafter. These simulations allowed us to compare the structural dynamics of the WT and P22L forms.

#### Umbrella sampling simulations of the Ape1 propeptide-propeptide interactions

To quantify the interaction strength between Ape1 propeptides for the WT and the P22L, we used umbrella sampling simulations to calculate the potential of mean force (PMF) as a function of the centre-of-mass distance between the two propeptides. AlphaFold models consistently predicted interaction interfaces within the first ∼20–24 N-terminal residues; therefore, we truncated each propeptide to its first 30 amino acids. Initial configurations were generated using steered MD pulling simulations using constant forces of 100, 50, and 20 kJ mol⁻¹ nm⁻². For the umbrella sampling, harmonic biasing potentials were applied with two force constants: 5000 kJ mol⁻¹ nm⁻² for separations < 2 nm (window spacing: 0.05 nm) and 1000 kJ mol⁻¹ nm⁻² for distances ≥ 2 nm (spacing: 0.1 nm). Each umbrella window was simulated for 100 ns, discarding the first 10 ns for equilibration. PMFs were reconstructed using the Weighted Histogram Analysis Method (WHAM)^84^ with a Jacobian correction applied as described previously^85^.

#### Triplet angle sampling from a disordered (High-T) LJ-fluid

To approximate the random triplet angle distribution, we simulated the LJ-fluid of argon, initializing 13^3^ atoms in a cubic lattice (11.3378^3^ nm^3^ box size) using a custom python script. MD simulations were run in the OPLS-AA force field (σ = 0.41 nm, ε = 0.99774 kJ mol⁻¹) for 100 ps in the NVT ensemble at 2000 K, a temperature chosen to generate a maximally disordered fluid rather than to model physical argon behaviour, and repeated a hundred times^86^. Triplet angle distributions were derived from all frames after the steady state was reached, with histograms averaged per-bin across replicates to yield mean and standard deviation profiles. Contact triplets were defined as having all three pair distances below d = 0.61 nm ≈ 1.5σ.

### Coarse-grained molecular dynamics simulations

We performed coarse-grained MD simulations to model the Ape1 condensates and access longer spatiotemporal timescales. We developed a patchy particle model to represent each Ape1 dodecamer^87,88^. An Ape1 was represented by a sphere with 12 interaction sites. These sites were positioned along the edges of a tetrahedron whose midsphere is the Ape1 dodecamer representation sphere, with two sites per edge, capturing the tetrahedral symmetry of the Ape1 dodecamer and its interacting propeptides. The interaction sites were separated 0.62 times their distance to the centre particle to approximate the expected distancee between propeptide exit sites on the edges of the Ape1 dodecamer.

#### Simulation setup

All coarse-grained simulations were performed using the software package LAMPPS (Large-scale atomic/ molecular massively parallel simulator)^89^ and were conducted using reduced units. The systems were thermalized with a Langevin thermostat^90^ (at kBT = 1) with a with a damping coefficient of 5τ. A time step of 0.005τ was used, where τ is the characteristic time scale is defined as τ= *^mσ2^ for all the particles, and σ=7.40 nm, is the radius of the Ape1 circumsphere. Particles interacted via the Lennard-Jones (LJ), defined as:

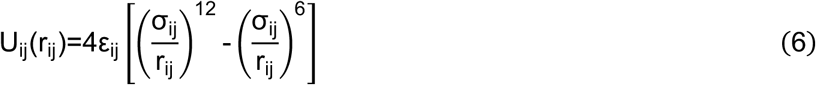

Where 𝑟_!"_ is the distance between two particles, σ_ij_ and ε_ij_ are the interaction length and energy scales between bead types ij. The values of ε_ij_ are normalized by kBT, the thermal energy. Our model has two types of particles: a type 1 central particle and type 2 interaction sites (Fig. 5a). We fixed ε_11_=0.1 and σ_11_=1.78 for steric repulsion, and optimize σ_22_and ε_22_ for the interaction sites to match experimental properties of the condensates.

#### Parameter optimization

To optimize the simulation parameters, we created an initial system of 343 (7x7x7) Ape1 patchy particles uniformly distributed in a cubic box simulation of size 42σ^3^. Using this system, we performed a grid search across a range of ε_22_∈ [0.5, 3.5] and σ_22_∈ [0.35, 0.9]. Each simulation ran for 5×10^7^ timesteps saving frames every 10^4^ timesteps. The last 500 frames were used for analysis. For each simulation, we determined the densities of the dense phase and the dilute phases, as well as the coordination number (*CN*). Parameters were selected to minimize the following objective function:

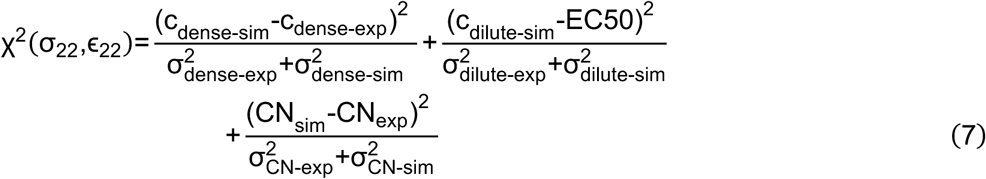

Here 𝑐_!_ are the dense and dilute concentrations in simulations and experiments. EC50 values were obtained from the spectral-shift dose response curves (Fig. 4c). The following values were obtained from the best picked tomograms (Fig. S9e): c_dense-exp_±σ_dense-exp_=451±49 μM and CN_exp_±σ_CN-exp_=12±4. The dense concentrations were determined from simulations using the k-nearest neighbour (kNN) algorithm^91^, similar to the description above for local concentration. This optimization yielded two distinct sets of parameters: one for the WT and another for the P22L mutant, with the latter corresponding to an increased interaction strength (Fig. S10a-d). Using these parameters, we performed production simulations with 1000 patchy particles (Ape1) in a cubic box (60𝜎^#^) for 1×10^8^ timesteps (Fig. 5b).

### Protein expression in E. coli

StrepII-NT*tag-HRV3C-Ape1 and StrepII-NT*-HRV3C-Ape1P22L fusion constructs (see Supplementary Table S2) were expressed from pET21a in E. coli BL21(DE3). MBP-Ape1 and MBP-Ape1P22L fusion constructs were expressed from pET28 in E. coli BL21(DE3). Cells were grown in lysogeny broth (LB) medium supplemented with ampicillin at 37 °C to an OD600 of 0.7, the temperature was reduced to 16 °C and expression was induced by the addition of 1 mM isopropyl-β-D-1-thiogalactopyranoside (IPTG) for 18 h. For StrepII-tagged fusion proteins, cells were pelleted at 3000g for 15 min at room temperature, resuspended in buffer W (100 Tris-HCl, 150 mM NaCl and 1 mM EDTA, pH 8.0) supplemented with 5% glycerol, 1% Triton X-100, 1 mM phenylmethylsulphonyl fluoride (PMSF), 1 mM DTT and cOmplete protease inhibitor cocktail (Roche) and lysed by sonication on ice. Cell lysates were cleared by centrifugation at 16,000g for 10 min at 4°C then applied to a Strep-TactinXT 4Flow column (IBA Lifesciences). The column was washed with Buffer W (100 mM Tris-HCl, 150 mM NaCl and 1 mM EDTA, pH 8.0) and the protein was eluted with Buffer BXT (100 mM Tris-HCl, 150 mM NaCl, 50 mM biotin and 1 mM EDTA, pH 8.0). For MBP-tagged proteins, cells were pelleted at 3000g for 15 min at room temperature, resuspended in buffer MBP (20 mM Tris-HCl, 200 mM NaCl, 1 mM EDTA, pH 7.4) supplemented with 5% glycerol, 1% Triton X-100, 1 mM phenylmethylsulphonyl fluoride (PMSF), 1 mM DTT and cOmplete protease inhibitor cocktail (Roche) and lysed by sonication on ice. Cell lysates were cleared by centrifugation at 16,000g for 10 min at 4 °C then applied to a MBPTrap column (Cytiva). The column was washed with Buffer MBP (20 mM Tris-HCl, 200 mM NaCl, 1 mM EDTA, pH 7.4) and the protein was eluted with Elution buffer (20 mM Tris-HCl, 200 mM NaCl, 1 mM EDTA, 10 mM Maltose, pH 7.4). All proteins were further purified over a high-load size exclusion Superdex200 column using 20 mM Hepes, 150 mM NaCl, pH 7.0. The purified protein was stored at −80 °C.

### In vitro Ape1 droplet formation

For unlabelled condensate formation, 10 µM of StrepII-NT*tag-HRV3C-Ape1 or StrepII-NT*- HRV3C-Ape1P22L of purified protein was incubated in 20 mM HEPES pH 7.0, 150 mM NaCl, and 5 mM DTT for 3 h and supplemented with 1 µL of NT*-HRV 3C protease (3.5 mg ml^-1^) and transferred into a Channel µ-Slide VI 0.4 (Ibidi). For labelled condensates, the reaction was additionally supplemented with 500 nM of StrepII-NT*tag-HRV3C-Ape1NT-647 or StrepII-NT*tag-HRV3C-Ape1P22LNT-647. All proteins were labelled according to manufacturer’s instructions using the Protein Labeling Kit RED-NHS 2^nd^ Generation (NanoTemper Technologies GmbH).

### In vitro FRAP analysis of Ape1 droplets

In vitro FRAP of Ape1 condensates was performed at room temperature using a DeltaVision OMX Super Resolution microscope operated by AcquireSR (4.5.10170-1). The image resolution was set to 256 × 256, and the excitation wavelength to 640 nm. For each sample, at least 13 frames were collected with a time interval of 15 s. Of those, one frame was collected before bleaching, and 12 frames were collected immediately after bleaching. Images were deconvolved using softWorX deconvolution plugin (version 7.2.1). Double and single normalisation was performed as described elsewhere^92^.

### Spectral shift analysis

MBP-Ape1 and MBP-Ape1^P22L^ were labelled as described previously^93^ using the Protein Labeling Kit RED-NHS 2nd generation (NanoTemper Technologies GmbH). Unreacted dye was removed in a gravity PD minitrap G25 column (Cytiva) equilibrated with the buffer containing 20 mM HEPES-KOH pH 7.5, 150 mM NaCl. A degree of labelling of 1.1-1.6 was typically achieved. The concentration of labelled MBP-Ape1 and MBP-Ape1^P22L^ was adjusted to 5 nM final concentration, with HEPES buffer supplemented with 0.01% Tween20. The ligand was submitted to 2-fold dilution series in HEPES buffer supplemented with 0.01% Tween20, producing a final ligand concentration ranging from 13 µM to 0.4 nM for MBP-Ape1 and, 6 µM to 0.2 nM for MBP-Ape1^P22L^. After 15 minutes incubation at room temperature, spectral shift (ratio 670/650 nm) was measured at 25°C with respective excitation powers of 10 and 20 % (WT) and 60 and 80 % (P22L) using a Monolith X (NanoTemper Technologies GmbH). Data from 3 (WT) and 4 (P22L) individual measurements were normalized using the MO.control software version 2.6.3 (NanoTemper Technologies GmbH). EC50 values were fitted with OriginPro 2020.

### Conformational ensemble of the Ape1 propeptide based on NMR chemical shifts

A conformational ensemble was derived from the Ape1 propeptide (residues 1-45 of the full-length protein) through an iterative protocol combining the statistical coil model generator flexible Meccano^94^ and the genetic algorithm ASTEROIDS^95^. Initially, a pool of 10,000 random coil conformers was generated, from which 200 conformations were selected to optimally match the experimental chemical shifts of CO, Cα, and Cβ nuclei that were previously published^14^. Using the Φ and ψ dihedral angles from these selected structures, a new library of 8500 conformers was constructed and combined with 1500 conformers from the original set. Another selection of 200 conformers was performed on this updated pool. This refinement cycle was carried out four times in total. Five independent ensembles of 200 conformers each were generated. Ensemble-averaged chemical shifts were predicted using SPARTA^96^, and secondary chemical shifts were computed with reference to RefDB^97^.

### Ape1 sequence homology search and multiple sequence alignment

Ape1 homologs were identified by performing a protein BLAST (blastp) search (https://blast.ncbi.nlm.nih.gov/Blast.cgi)^98^ of the full-length *S. cerevisiae* Ape1 sequence (Uniprot ID: P14904) against the ClusteredNR database (accessed 13/10/2025) using a BLOSUM62 matrix. From the 50 homologous sequences identified, the following 10 were used as inputs for multiple sequence alignment: *Saccharomyces arboricola* H-6 (sequence ID: EJS42965.1), *Naumovozyma dairenensis* CBS 421 (sequence ID: XP_003672705.1), *Maudiozyma saulgeensis* (sequence ID: SMN18868.1), *Torulaspora delbrueckii* (sequence ID: XP_003678797.1), *Zygosaccharomyces parabailii* (sequence ID: AQZ16088.1), *Lachancea nothofagi* CBS 11611 (sequence ID: SCU89774.1), *Kluyveromyces lactis* (sequence ID: QEU62558.1), *Candida maltosa* Xu316 (sequence ID: EMG50408.1), *Vanderwaltozyma polyspora* DSM 70294 (sequence ID: XP_001642941.1) and *Monosporozyma unispora* (sequence ID: KAG0661901.1). The species in which these sequences are found were selected as representative examples from diverse genera belonging to the taxonomic family Saccharomycetaceae. Multiple sequence alignment was performed using the Clustal Omega tool provided by the EMBL-EBI Job Dispatcher (https://www.ebi.ac.uk/jdispatcher/msa/clustalo)^99,100^ and the resulting alignment and consensus sequence of the propeptide region (Fig. S10f) were visualised using the ggmsa package implemented in R (http://yulab-smu.top/ggmsa).

### Statistics and reproducibility

For all cell biology experiments (Fig. 2e and 2f, Fig. S1b and S5b-f), at least 3 independent biological replicates were performed. For in all vitro experiments (Fig. 4c, 5c and 5d, Fig. S9h-i and S10g-m), at least 3 technical replicates were performed. The number of replicates performed for each experiment is stated in the corresponding figure legends. Statistical tests were performed using GraphPad Prism (version 10.5.0). Statistical significance was determined by two-tailed paired t-test or two-way ANOVA with Tukey or Šídák multiple comparisons test as specified in the figure legends. Data was assumed to be normally distributed but this was not formally tested. For live cell fluorescence microscopy, acquisition areas were chosen at random based on brightfield images and, where possible, image analysis was automatised with identification of fluorescence signal performed by intensity-based thresholding. Further quantification and statistical analysis details are included in the relevant Materials and Methods sections.

## Acknowledgements

We thank the members from the Mechanisms of Cellular Quality Control research group and the Central Electron Microscopy Facility at the Max Planck Institute of Biophysics, Anna Bieber, Cristina Capitanio, Florian Beck, Beata Turonova, Iskander Khusainov, Martin Beck, Andre Schwarz, the MPI Brain Research Imaging Facility and the staff at the Max Planck Computing and Data Facility for support and discussions. In particular, we thank Nobuo N. Noda (BIKAKEN) for providing access to the original published NMR data. We thank Martin Blackledge and Malene R. Jensen for analysis scripts and providing access to flexible meccano and ASTEROIDS. We thank Evgeny V. Mymrikov for support and discussion of spectral-shift analysis. The authors also acknowledge the use of the large language model GPT-4o (OpenAI) for writing initial code snippets and debugging errors. The work was supported by the Max Planck Society, the European Union (F.W.: ERC, IntrinsicReceptors, 101041982, C.K.: ERC, AutoClean, 769065, S.M.: ERC, MultiMotif, 802209), the Deutsche Forschungsgemeinschaft (F.W., C.K., G.H., SFB 1177; project ID 259130777, C.H., C.K. SFB1381 project ID 403222702, project IDs 450216812, and 409673687, under Germany’s Excellence Strategy CIBSS - EXC-2189 project ID 390939984), the Leibniz-Forschungsinstitut für Molekulare Pharmakologie (S.M.), the Chan Zuckerberg Initiative for Visual Proteomics Imaging (G.H., grant number 2021-234666), and the Clusterproject ENABLE funded by the Hessian Ministry for Science and the Arts (G.H.). Views and opinions expressed are, however, those of the author(s) only and do not necessarily reflect those of the European Union or the ERC Executive Agency. Neither the European Union nor the granting authority can be held responsible for them.

## Author Contributions

Conceptualization: E.B., S.C.L, J.L., J.B., C.K., G.H. and F.W.; Methodology: E.B., S.C.L, J.L., J.B., H.M., J.F.M.S., J.J.N., S.M., S.W., C.H., C.K., G.H. and F.W.; Software: E.B., S.C.L, J.L., J.B.; Validation: E.B., S.C.L, J.L., J.B., H.M., J.F.M.S., J.J.N., S.M., S.W., C.H., C.K., G.H. and F.W.; Investigation: E.B., S.C.L, J.L., J.B., H.M., J.F.M.S., J.J.N., S.M.; Writing – original draft: E.B., J.L., F.W.; Writing – review and editing: E.B., S.C.L, J.L., J.B., H.M., J.F.M.S., J.J.N., S.M., S.W., C.H., C.K., G.H. and F.W.; Supervision: S.W., C.H., C.K., G.H. and F.W.; Project administration: G.H. and F.W.; Funding acquisition: S.M., C.H., C.K., G.H. and F.W.

## Competing Interests

The authors declare no competing interests.

## Supplementary Materials

### Supplementary Figures

**Figure S1:**
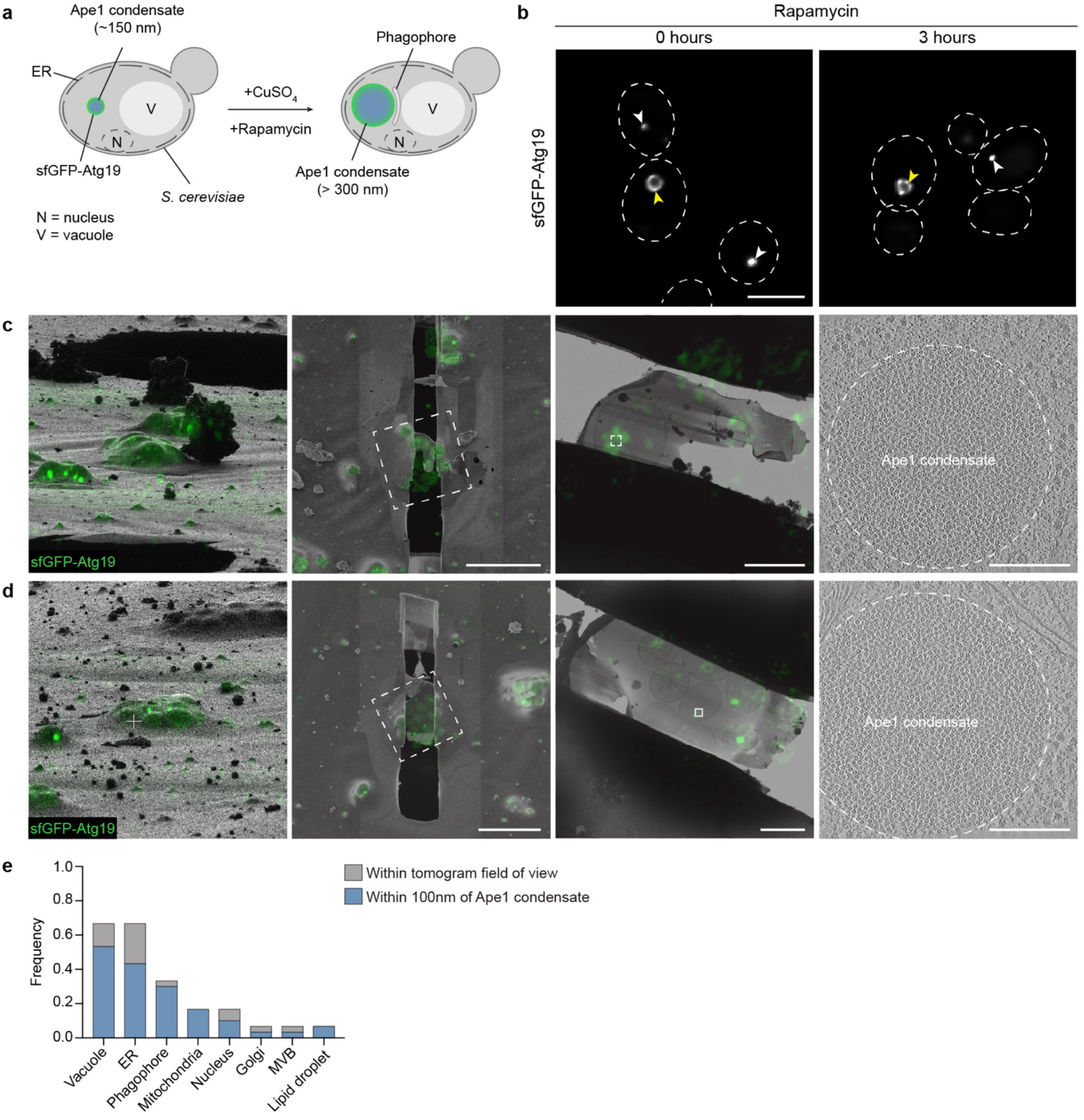
Correlative cryo-ET of Ape1 condensates in *S. cerevisiae*. **a)** Schematic of sample preparation prior to EM grid preparation. **b)** Representative deconvolved fluorescence microscopy images of sfGFP-Atg19 signal in cells overexpressing WT Ape1 before or after rapamycin treatment. sfGFP-Atg19 signal is visible as rings (yellow arrows) and puncta (white arrows). Scale bar = 5 µm. **(c-d)** Additional examples of correlation workflow as shown in Fig. 1b). Maximum intensity projections of sfGFP-Atg19 signal are overlaid on FIB images of the targeted lamella site, SEM images of the milled lamellae and TEM images of the milled lamellae at the regions indicated by the white boxes on the SEM lamella images. Scale bars = 5 µm. Central slices of denoised tomograms reconstructed from tilt series taken at the positions indicated by the white boxes on the TEM lamella images are shown. Scale bars = 250 nm. **e)** Quantification of frequency of tomograms containing the selected organelles within the same field of view as an Ape1 condensate (grey) or within 100 nm of the Ape1 condensate (blue). Bars are overlaid.

**Figure S2:**
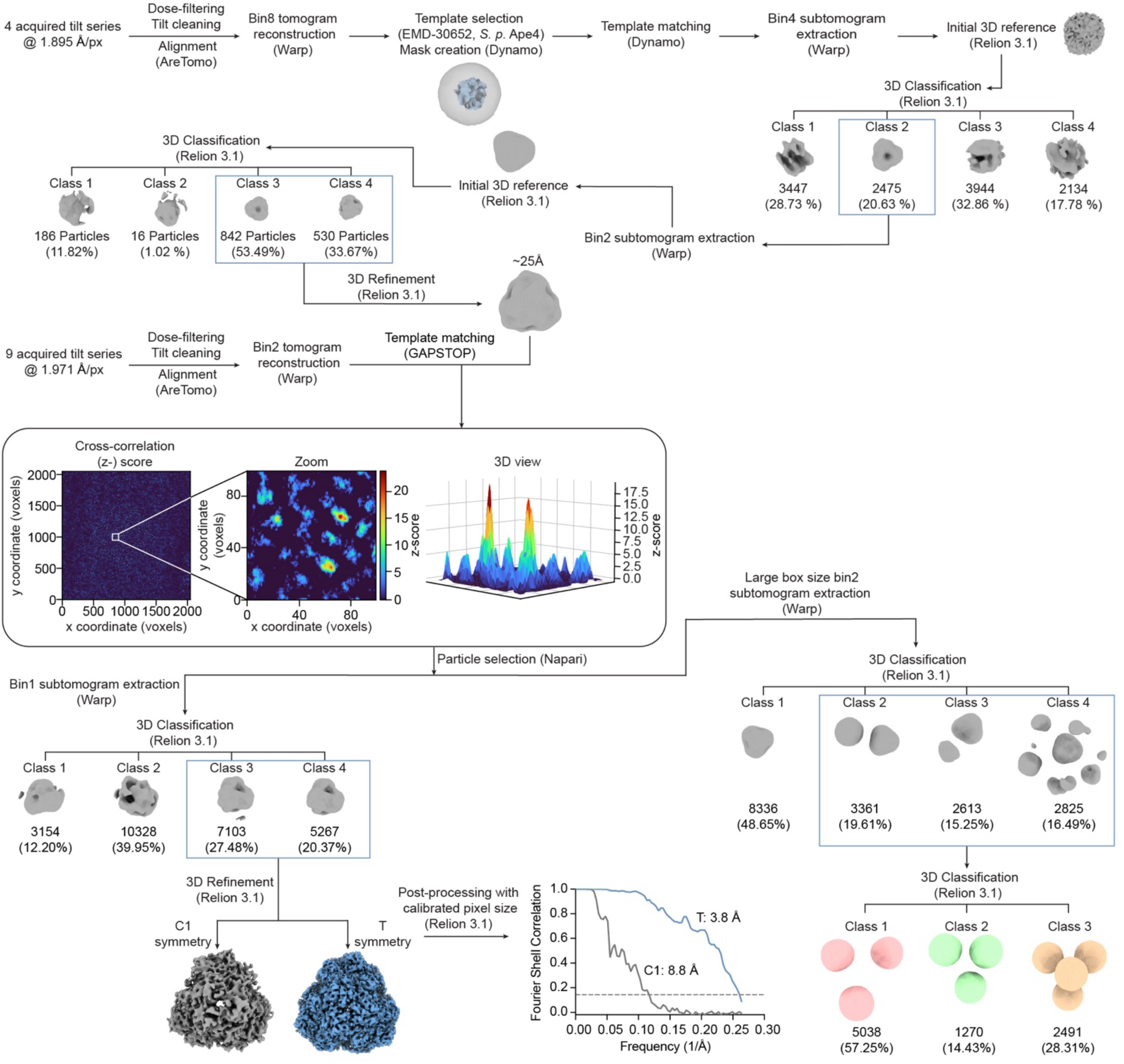
Summary of the image processing pipeline applied for the Ape1 dodecamer. Tomograms were reconstructed at bin8 for initial inspection. Initial template matching was performed in Dynamo using the EMDB entry of structural homologue Ape4 from *S. pombe* (EMDB: EMD-30652) as a reference^28^. Coordinates were extracted in Warp and RELION 3.1 was used for 3D classification and refinement of an initial in situ data-derived Ape1 dodecamer reference. A second round of template matching was performed with this reference at bin2 using GAPSTOP^TM^. Subtomograms were extracted around the coordinates of the cross-correlation peaks using Warp, and were then used for 3D classification and refinement to clean particle lists in RELION. FSC curves corresponding to the refinements of the selected particles using C1 (grey) and tetrahedral symmetry (blue) are shown (threshold line at FSC = 0.143). Averages containing several neighbouring Ape1 dodecamers calculated following extraction of particles using a larger box are shown in orange, pink and green. Numbers of particles are shown for each step under their corresponding class.

**Figure S3:**
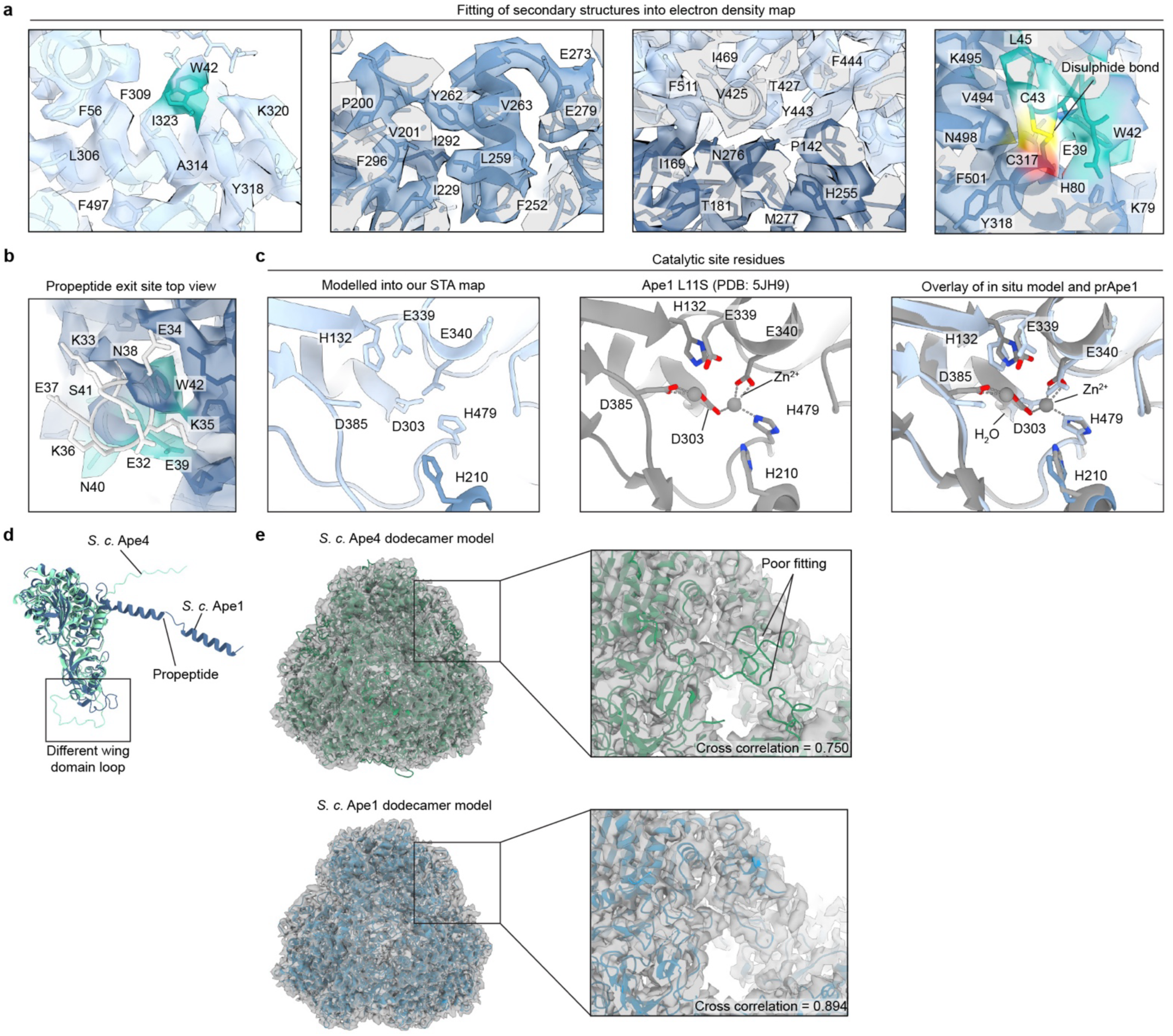
High resolution features of the in situ Ape1 dodecamer structure. **a)** Fitting of secondary structure elements across different regions of the in situ Ape1 map. **b)** Top view of the N-terminal propeptide exit site showing the in situ Ape1 map and fitted atomic model. Residues unable to be modelled into the electron density map are coloured in grey and the fitted residues and corresponding electron density are shown in cyan. **c)** Comparison of the Ape1 catalytic residues between the in situ model (blue) and the previously available crystal structure (PDB: 5JH9, grey)^14^. **d)** Comparison of the AlphaFold predictions of monomeric *S.c.* Ape4 (green) and *S.c.* Ape1 (blue). **e)** Top: fitting of an *S.c.* Ape4 dodecamer generated with SWISS-MODEL (green) into the electron density map. Bottom: fitting of the in situ Ape1 dodecamer model (blue) into the map. Map-in-map cross-correlation values calculated using the ChimeraX fitmap command optimising for correlation are indicated.

**Figure S4:**
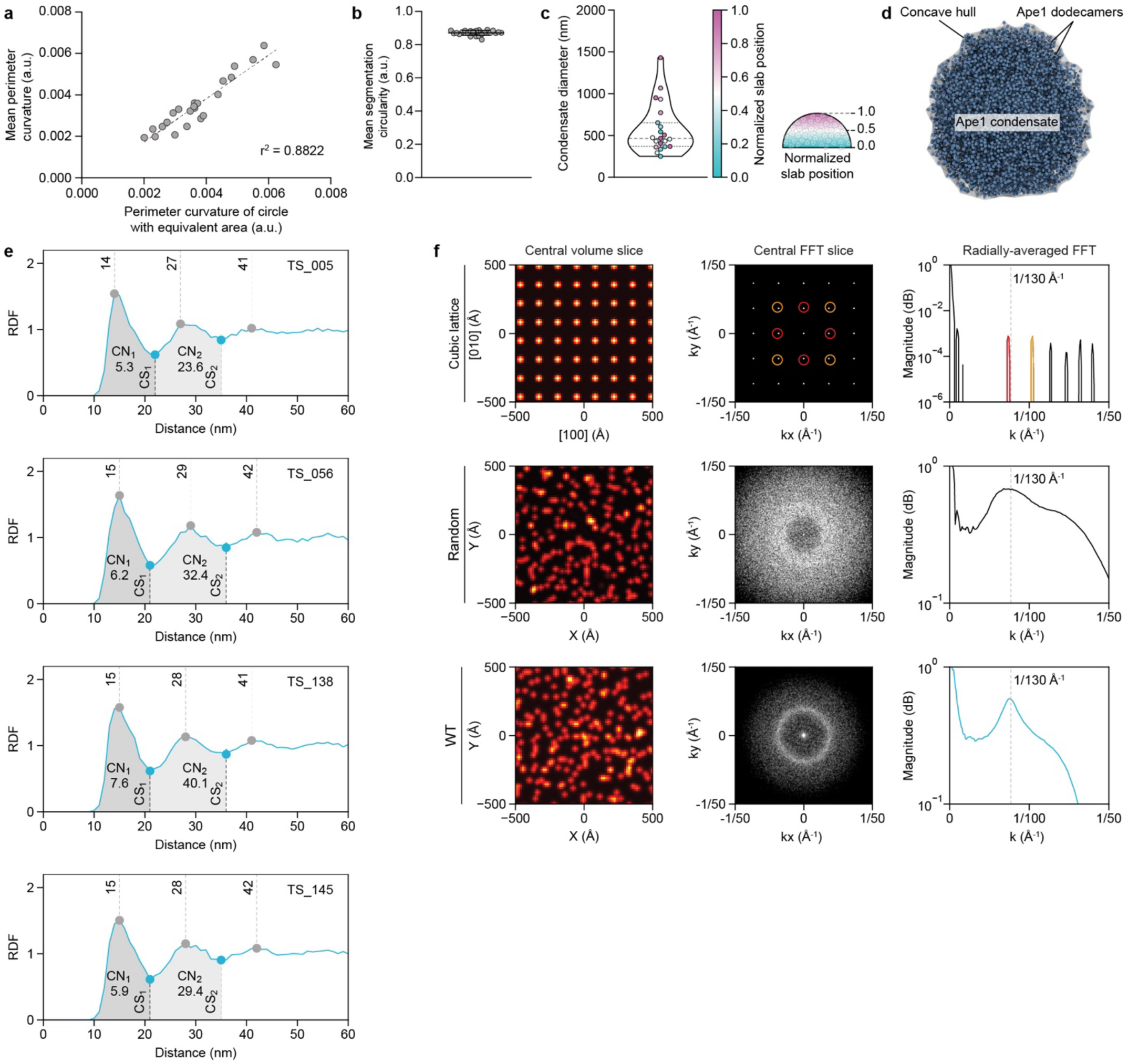
Analysis of Ape1 condensate properties from tomographic data. **a)** Correlation between mean perimeter curvature of segmented central slices of Ape1 condensates analysed in Fig. 2c) and perimeter curvature of circles of equivalent area. Dashed line is the best-fit line calculated by simple linear regression. **b)** Mean circularity of segmentations of 3 single slices of each of the Ape1 condensates analysed in Fig. 2c). Error bars show mean ± standard deviation. **c)** Violin plot of estimated Ape1 condensate diameter based on sphere fitting. Individual data points indicate individual Ape1 condensates and dashed lines indicate the median diameter and 25 and 75% quartiles. Colour indicates the normalised slab position i.e. distance of the tomogram slice from the pole of the Ape1 condensate. **d)** Concave hull enclosing the Ape1 dodecamer particle positions for a representative WT tomogram. **e)** RDFs of individual WT Ape1 condensates contributing to the average profile shown in Fig. 2d). Dashed lines show successive neighbour shells with their corresponding distances and shaded regions delineate CS1 and CS2 with stated coordination numbers (CN1 and CN2). **f)** Averaged, log-normalised 3D FFT of particle distributions. Top row: Simple-cubic lattice with a 13 nm lattice constant denoted in Miller notation. Middle row: Poisson-disk packing of 13 nm-diameter spheres. Bottom row: Ape1 particle coordinates from a representative WT tomogram. Left panels show summed-intensity projections from a central slab of the simulated volume. Centre panels depict the central slice of the enlarged and histogram-adjusted FFT. Right panels plot the 3D radial average; for the WT Ape1 particles, the local maximum at 1/130 Å^-1^ is highlighted and for the cubic lattice, the corresponding diffraction spots in the central FFT slice are colour-coded.

**Figure S5:**
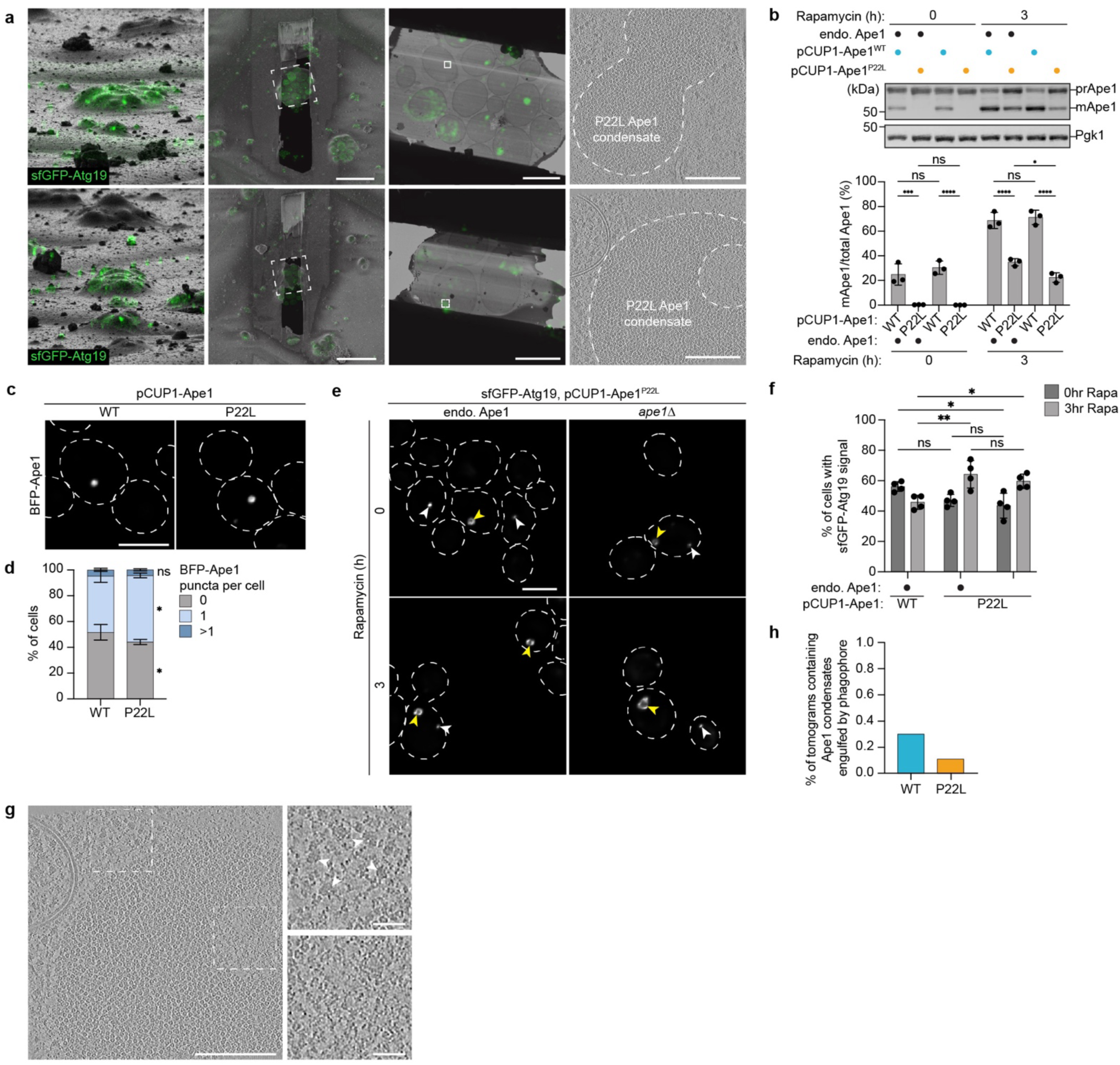
Correlative cryo-ET of P22L Ape1 condensates in *S. cerevisiae*. **a)** Examples of correlation workflow used to acquire tilt series of Ape1 condensates containing P22L Ape1 dodecamers. Maximum intensity projections of sfGFP-Atg19 signal are overlaid on FIB images of the targeted lamella site, SEM images of the milled lamellae and TEM images of the milled lamellae at the regions indicated by the white boxes on the SEM lamella images. Scale bars = 5 µm. Central slices of denoised tomograms reconstructed from tilt series taken at the positions indicated by the white boxes on the TEM lamella images are shown. Scale bar = 250 nm. **b)** Immunoblotting of Ape1 in cells with or without endogenous levels of WT Ape1 and additionally overexpressed WT or P22L Ape1 before and after treatment with rapamycin. Pgk1 is used as a loading control. For quantification of Ape1 processing, error bars represent the mean ± standard deviation. Statistical significance was determined by two-way ANOVA with Tukey multiple comparisons test. ns P > 0.05, * P < 0.05, ** P < 0.01, *** P < 0.001, **** P < 0.0001 (P = multiplicity adjusted P value). n = 3 biological replicates. **c)** Representative deconvolved fluorescence microscopy images of BFP-tagged endogenous WT Ape1 signal in cells containing overexpressed untagged WT or P22L Ape1. Scale bar = 5 µm. **d)** Quantification of proportion of cells containing Ape1 puncta. Error bars represent mean ± standard deviation. Statistical significance was determined by two-way ANOVA with Šídák multiple comparisons test. ns P > 0.05, * P < 0.05, ** P < 0.01, *** P < 0.001, **** P < 0.0001 (P = multiplicity adjusted P value). n = 4 biological replicates. **e)** Representative deconvolved fluorescence microscopy images of sfGFP-Atg19 signal in cells with or without endogenous levels of WT Ape1 and additionally overexpressed P22L Ape1 before and after treatment with rapamycin. sfGFP-Atg19 signal is visible as rings (yellow arrows) and puncta (white arrows). Scale bar = 5 µm. **f)** Quantification of proportion of cells containing localised Atg19 signal. Error bars represent mean ± standard deviation. Statistical significance was determined by two-way ANOVA with Tukey multiple comparisons test. * P < 0.05, ** P < 0.01, *** P < 0.001, **** P < 0.0001 (P = multiplicity adjusted P value). n = 4 biological replicates. **g)** Denoised central tomogram slice of P22L Ape1 condensate with enlarged cytosolic (top) and hole (bottom) regions. White boxes indicate enlarged regions. White arrows indicate ribosomes. Scale bars = 250nm (full tomogram slice), 50nm (insets). **h)** Quantification of percentage of tomograms containing Ape1 condensates being engulfed by phagophore membranes.

**Figure S6:**
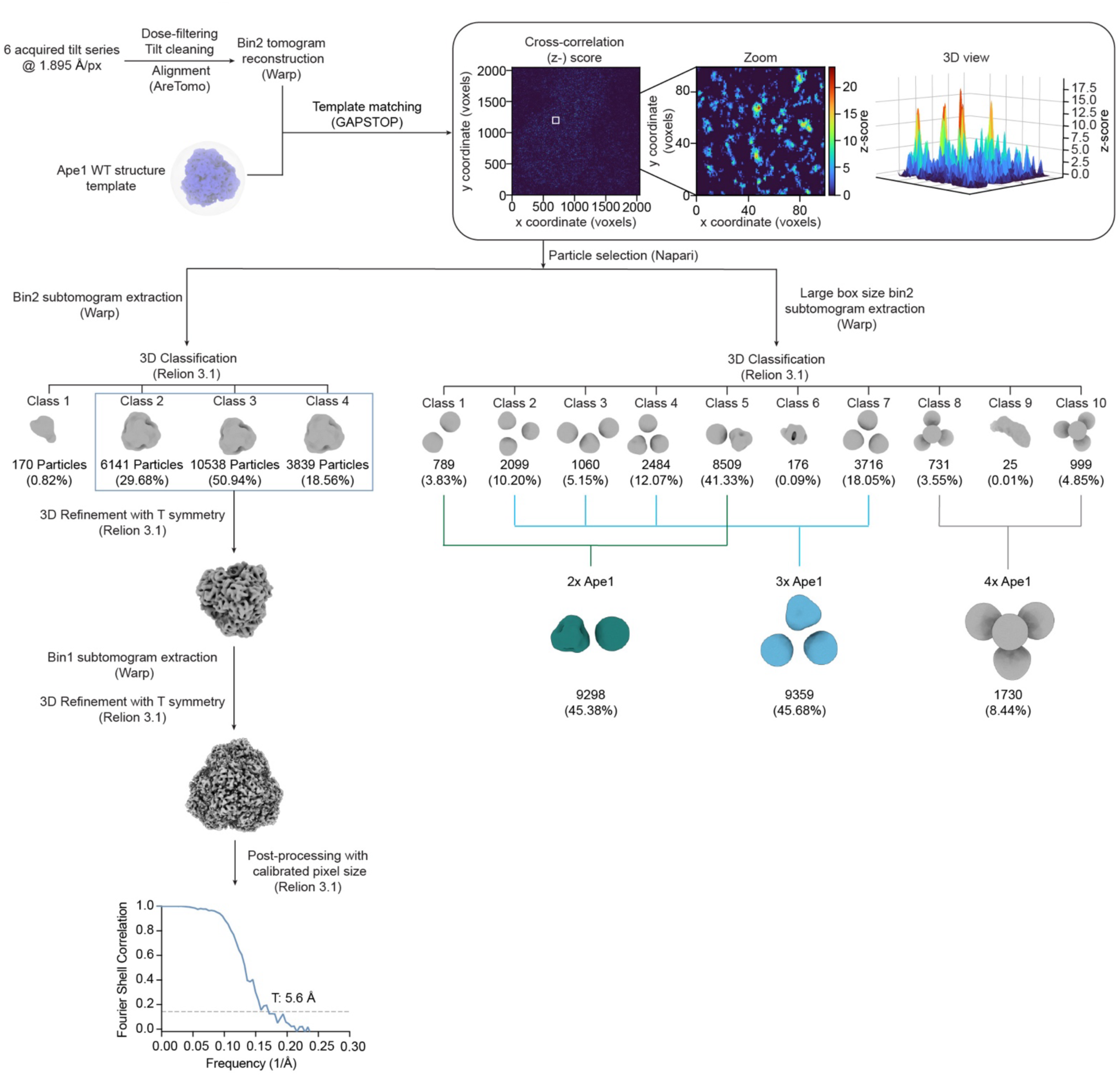
Summary of the image processing pipeline for Ape1 dodecamers in the presence of endogenous WT and overexpressed P22L Ape1. Tomograms were reconstructed at bin8 for initial inspection. Template matching was performed at bin2 with GAPSTOP^TM^ using the Ape1 WT average as a reference. Subtomograms around the coordinates of the cross-correlation peaks were extracted in Warp and 3D classification in RELION 3.1 was used to clean particle coordinates. FSC curve corresponding to the refinement of the selected particles imposing tetrahedral symmetry, yielding a map of 5.1 Å resolution, is shown (threshold line at FSC = 0.143). Averages containing several neighbouring Ape1 dodecamers calculated following extraction of particles using a larger box are shown in teal, cyan and grey. Numbers of particles are shown for each step under their corresponding class.

**Figure S7:**
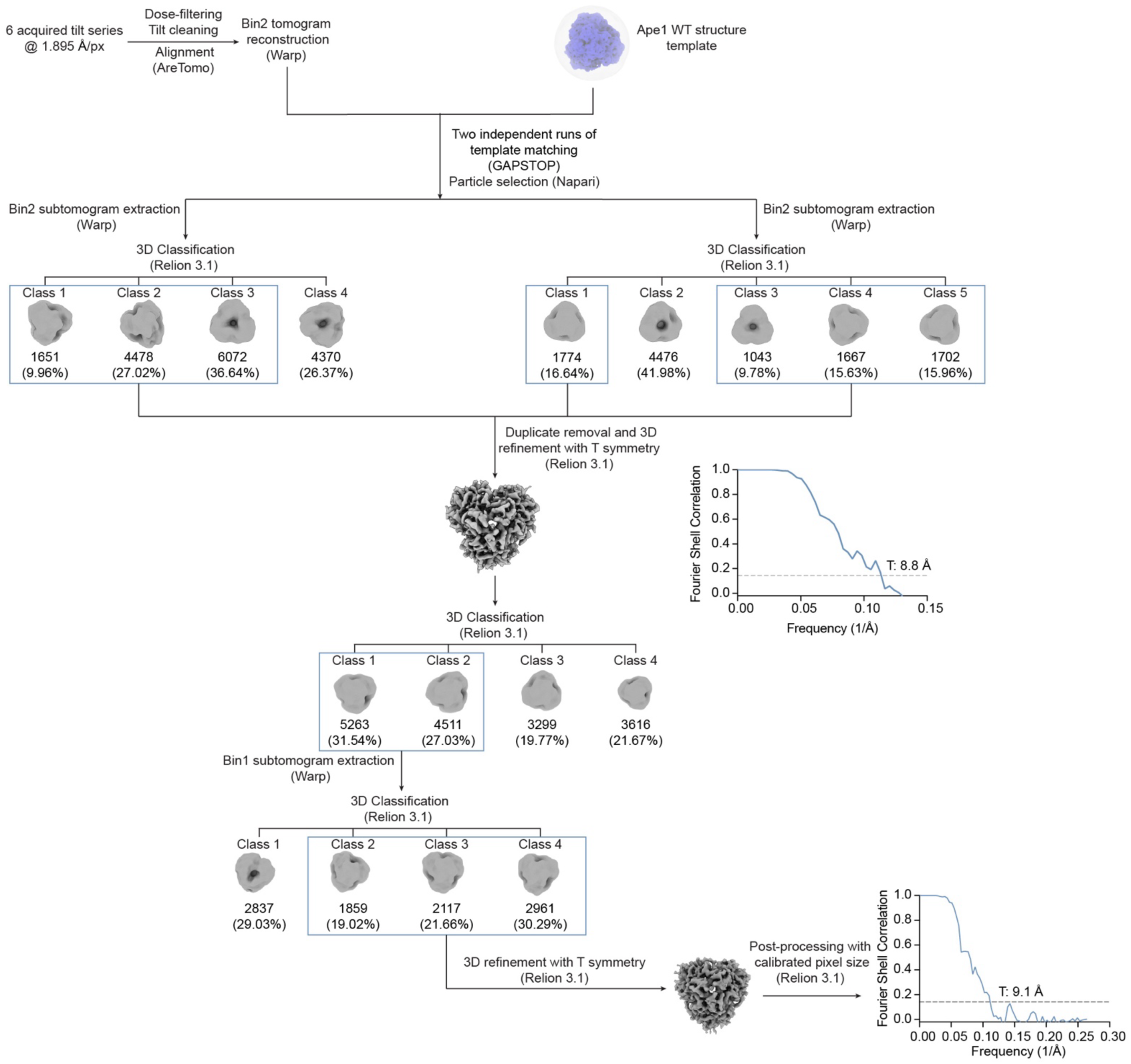
Summary of the image processing pipeline for Ape1 P22L dodecamers in the absence of WT Ape1. Tomograms were reconstructed at bin8 for initial inspection. Template matching was performed at bin2 with GAPSTOP^TM^ using the Ape1 WT average as a reference. Subtomograms around the coordinates of the cross-correlation peaks were extracted in Warp. Subtomograms from two independent acquisitions were used for 3D classification to clean particle lists for Ape1 classes. Then particles were merged and refinement using T symmetry was performed. After this, one more round of 3D classification was performed and subtomograms were extracted at bin1, further classified and refined. Numbers of particles are shown for each step under their corresponding class. FSC curve corresponding to the refinement of the selected particles imposing tetrahedral symmetry is shown (threshold line at FSC = 0.143).

**Figure S8:**
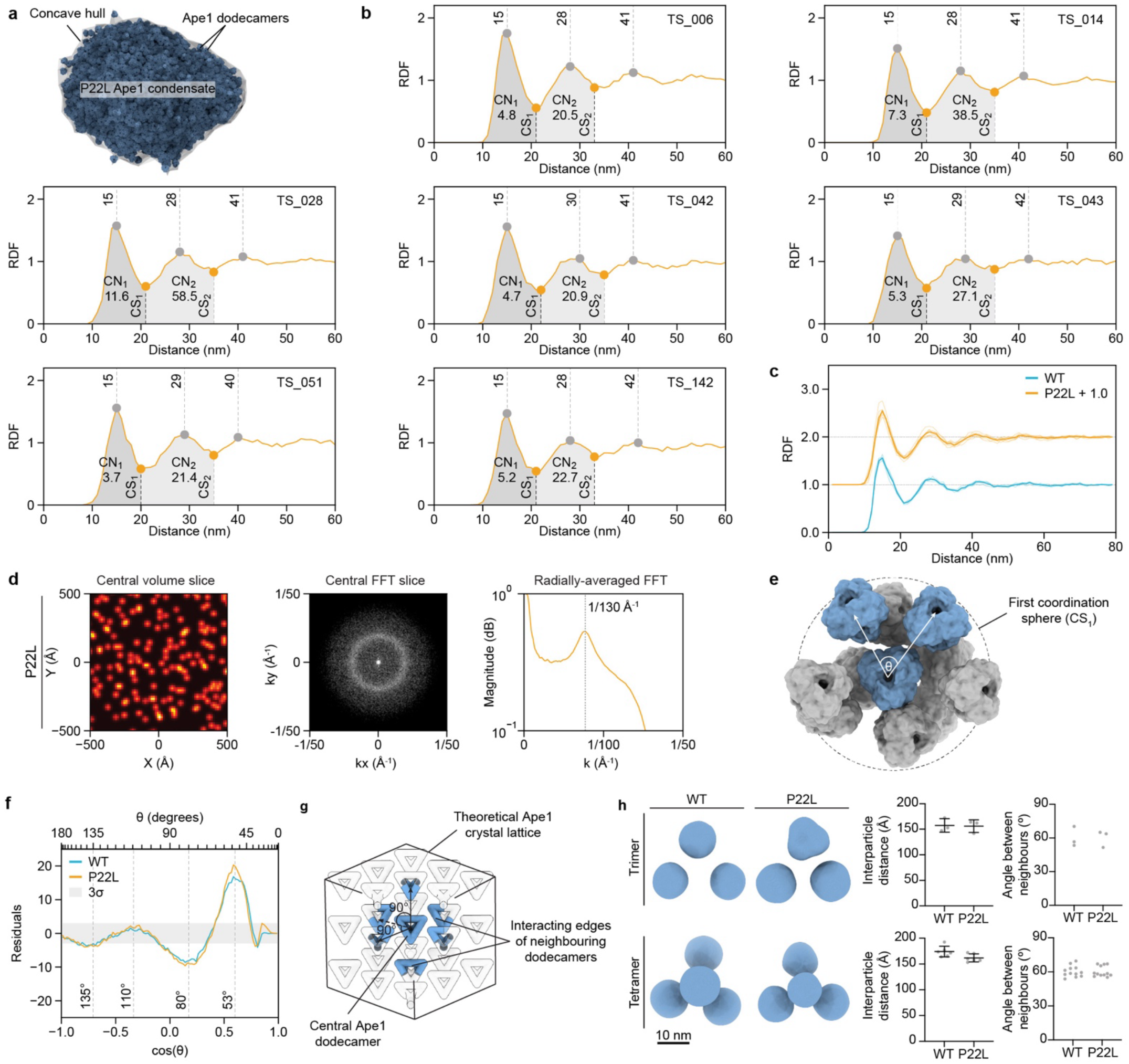
Local organisation of WT and P22L Ape1 condensates is indistinguishable at the level of local interparticle distances and angles. **a)** Concave hull enclosing the particle positions for a representative P22L tomogram. **b)** RDFs of individual P22L Ape1 condensates contributing to the average profile shown in Fig. 3c). Dashed lines show successive neighbour shells with their corresponding distances and shaded regions delineate CS1 and CS2 with stated coordination numbers (CN1 and CN2). **c)** RDFs of Ape1 condensates containing overexpressed WT or P22L Ape1. Solid lines show the average profile calculated from n = 4 WT (blue) and n = 7 P22L (orange) tomograms and transparent lines indicate the profiles for individual tomograms. For visualization purposes, P22L line profiles have been shifted in y by + 1.0. **d)** Averaged, log-normalised 3D FFT of Ape1 particle coordinates for all P22L tomograms. Left panel shows a summed-intensity projection from a central slab of the simulated volume. Centre panel displays the 3D FFT, enlarged and histogram-adjusted around the central ring. Right panel presents the 3D radial average, highlighting the local maximum at ∼1/130 Å⁻¹. **e)** Central Ape1 dodecamer and its neighbours within CS1. Particles shown in blue represent the particles between which vectors are calculated to determine the triplet angle (θ). **f)** Standardised residuals of angular distribution between WT (blue) or P22L (orange) Ape1 condensates and a random angular distribution as shown in Fig. 3e). The grey area indicates 3 standard deviations of the random distribution. **g)** Schematic of theoretical cubic lattice of Ape1 dodecamers (represented by tetrahedra) formed by interactions between propeptides protruding from perpendicular edges of neighbouring dodecamers. A single central tetrahedron and the interacting edges from 6 neighbouring dodecamers are shown in blue. **h)** Selected 3D classes of 3 or 4 neighbouring Ape1 dodecamers from the WT and P22L Ape1 condensates (see Fig. S2 and S6). Scale bar = 10 nm. Graphs indicate the measured interparticle distances and inter-neighbour vector angles within each class. Error bars represent the mean ± standard deviation.

**Figure S9:**
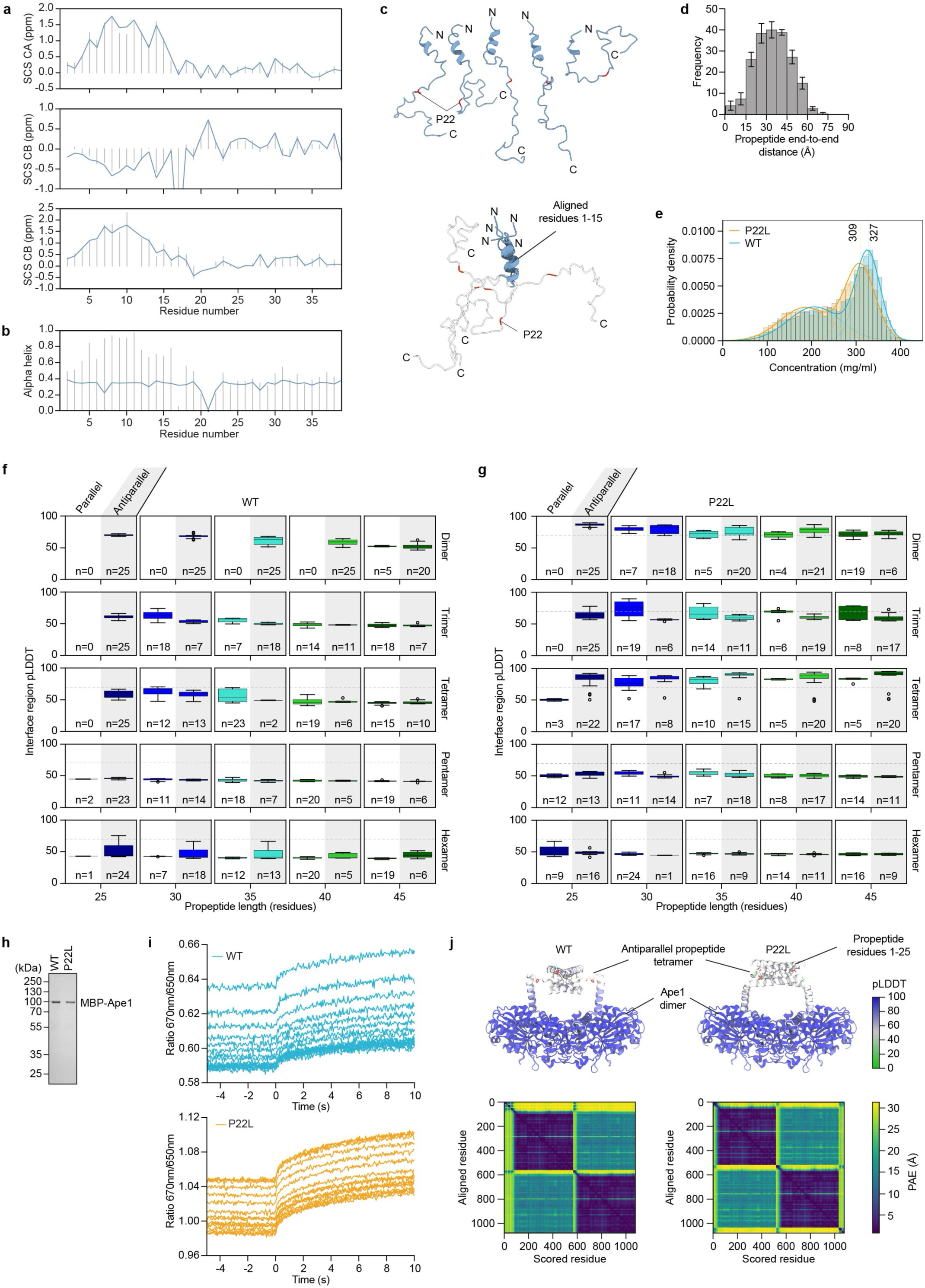
Characterisation of the Ape1 propeptide in isolation and upon binding. **a)** Back calculation of substituent chemical shifts (SCSs) (Calpha – CA, Cbeta – CB, CO) from a conformational ensemble derived from a combination of flexible meccano and ASTEROIDS is shown as blue lines above experimental values from previous acquired NMR data in grey^14^. **b)** Helical content of propeptide conformers. Grey bars represent the sampling of the conformational ensemble derived from the chemical shifts^14^, blue lines represent the sampling of a statistical coil (flexible meccano) ensemble. Alpha helical region is defined as ϕ < 0°, −120° < ψ < 50° ^47^. **c)** Individual conformers of propeptide residues 1-45 selected from a conformational ensemble that was generated based on NMR chemical shifts^14^ using a combination of the statistical coil generator flexible meccano and the genetic algorithm ASTEROIDS. Conformers are also shown aligned between residues 1-15. Aligned residues are shown in blue with the remainder of the propeptide shown in grey. **d)** Frequency of average propeptide end-to-end distances for each of 5 ensembles containing 200 conformers each, calculated based on NMR chemical shifts. Column heights represent the mean frequency of conformers for each distance bin. Error bars represent the standard deviation. **e)** Concentration of Ape1 dodecamers within each Ape1 condensate calculated using the local density distribution of the best picked tomograms for WT (blue) and P22L (orange). Dashed curved lines indicate the fitted Gaussian functions from a Gaussian-Mixture-Model. Dashed vertical lines show the mean of the largest Gaussian with their respective concentration value. **f)** Interface (residues 1 to 25 with a heavy atom within 5 Å of a heavy atom of residues 1 to 25 of a different chain) pLDDT scores for AlphaFold 2.3 predictions of different WT Ape1 propeptide oligomers. The length of the propeptide was varied from 25 to 45 residues in 5 residue steps. Plots show scores for 25 predictions per length sorted by parallel or antiparallel arrangement of the N-terminal propeptide helices. The absolute number of predicted structures displaying the respective orientation is listed below each box-and-whiskers plot. Boxes indicate interquartile range and the central line represents the median. Whiskers in each direction extend to the furthest data point within 1.5x(interquartile range) from the edge of the box. Points indicate outliers beyond that range. The grey dashed line indicates a pLDDT score of 70. **g)** Same as Fig. S9f) but for P22L propeptides. **h)** Purification of MPB-tagged WT and P22L Ape1 monomers for spectral-shift analysis. **i)** Fluorescence raw traces of spectral-shift dose response analysis shown in Fig. 4c). **j)** Left: AlphaFold 3 prediction of 2 WT full-length Ape1 monomers and 2 WT propeptide fragments (residue 1-25) with the same propeptide interface as the AlphaFold 2.3 predicted tetramer in Fig. 4a). Coloured by pLDDT. Sidechains of Leu, Val, Ile, and Pro residues in the propeptide are shown as sticks. Selected from 5 models, also top scoring model. Plot below shows predicted aligned error (PAE), where lower values indicate a higher confidence in the relative placement of two residues with respect to each other. Right: AlphaFold 3 prediction of 2 P22L full-length Ape1 monomers and 2 P22L propeptide fragments (residue 1-25) with the same propeptide interface as the AlphaFold 2.3 predicted tetramer in Fig. 4a). Coloured by pLDDT. Sidechains of Leu, Val, Ile, and Pro residues in the propeptide are shown. Selected from 550 models, global score close to top scoring model. Plot below shows PAE. Note that the order of protein chains in the PAE plots is different in WT and P22L predictions: in WT, the two propeptide chains come before the two full-length chains, in P22L, the order is reversed.

**Figure S10:**
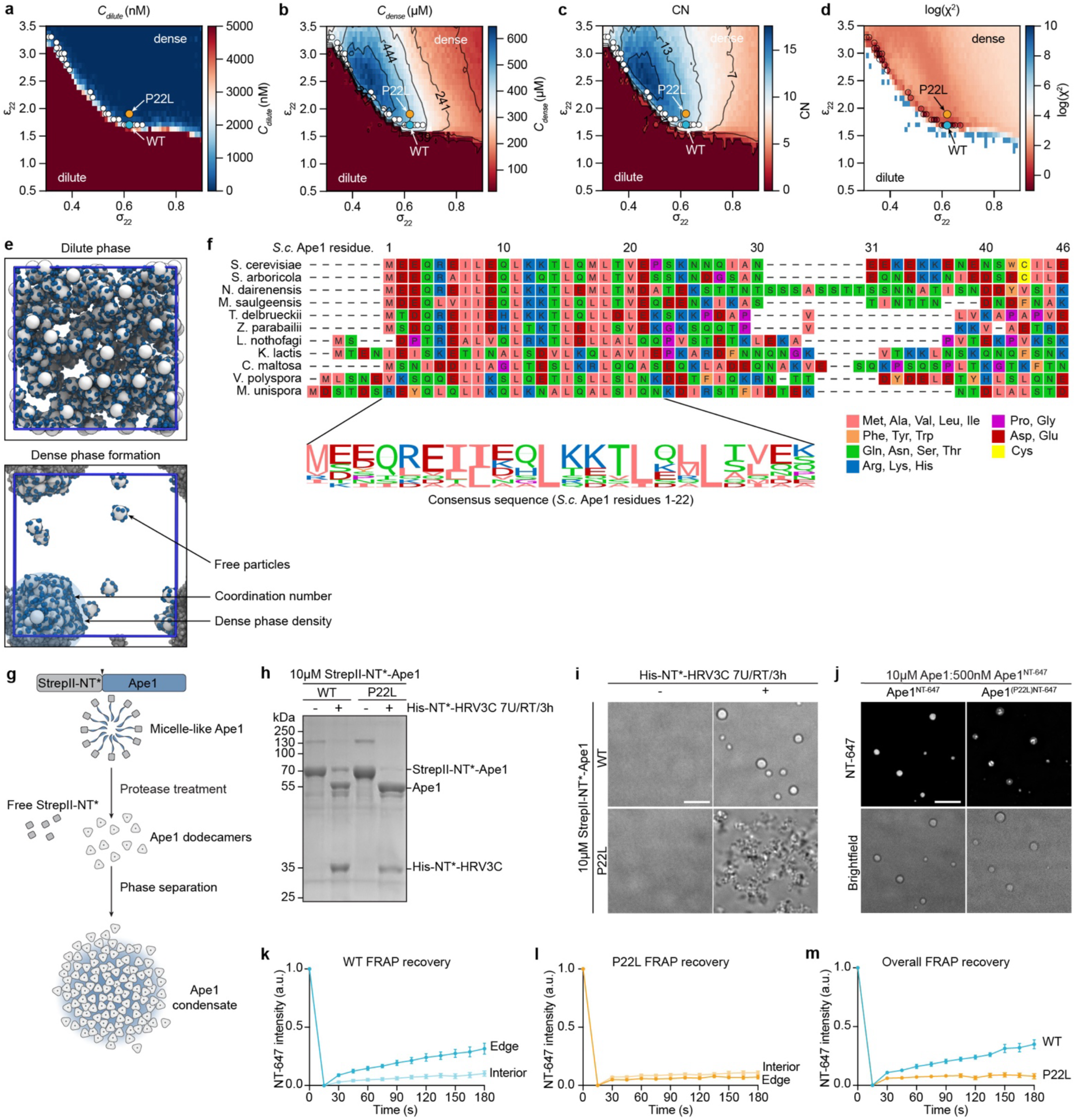
In vitro reconstitution and coarse-grained simulations confirm differences between WT and P22L Ape1 condensate viscosity. a-d) Optimisation of the ε_22_ and σ_22_ parameters of the model (see Materials and Methods eq. 6). The panels show (A) the dilute (c_dilute_) and (B) dense (c_dense_) concentrations and (C) the average number of neighbours (*CN*). The interaction parameters were sampled on a grid (ε_22_∈ [0.5, 3.5] and σ_22_∈ [0.35, 0.9]) and the optimal values, shown as a blue (WT) and orange (P22L) circles, were obtained by minimizing χ^2^ (see Materials and Methods eq. 7). **e)** Snapshots of coarse-grained simulations showing the effect of varying ε_22_ and σ_22_. The simulations produce a dilute phase (top), a dense + dilute phase (bottom) or only dense phase (not shown). **f)** Clustal multiple sequence alignment of selected Ape1 sequence homologs from various budding yeast genera identified by BLAST search using the full-length *S. cerevisiae* Ape1 sequence. Residues are coloured according to their physicochemical properties and the alignment shows all sequences from their N-terminus up to and including the *S. cerevisiae* propeptide sequence residue 46. The consensus sequence for amino acids corresponding to residues 1-22 of the *S. cerevisiae* sequence is shown below the alignment. **g)** Schematic of in vitro Ape1 condensate reconstitution. Ape1 was N-terminally tagged with StrepII-NT* to enhance solubility by forming micelle-like particles during purification. Upon removal of the StrepII-NT* tag, Ape1 spontaneously forms protein condensates. 10 µM of recombinant StrepII-NT*-Ape1WT and StrepII-NT*-Ape1P22L were subjected to protease cleavage to remove the solubility tag. **h)** SDS-PAGE gel of purified StrepII-NT*-Ape1 in the presence or absence of protease treatment (His-NT*-HRV3C) demonstrating the efficiency of tag cleavage. **i)** Brightfield microscopy images of structures formed by StrepII-NT*-Ape1 WT or P22L ± protease treatment. Scale bar = 5µm. **j)** Fluorescent Ape1 condensates were formed mixing 10 µM of unlabelled StrepII-NT*-Ape1WT and 500 nM of NT-647 dye-labelled StrepII-NT*-Ape1WT or StrepII-NT*-Ape1P22L. Samples were subjected to protease cleavage to remove the solubility tag and analysed by fluorescence microscopy. Scale bar = 5 µm. **k-m)** Quantification of fluorescence recovery after photobleaching on reconstituted Ape1 droplets. Recovery was measured at edges and interior of the droplets for WT (k) and P22L (l). The overall recovery was compared between WT and P22L (m). Error bars represent the mean ± standard error of the mean for n = 10 structures per condition per replicate. 3 technical replicates were performed for each condition.

### Supplementary Tables

**Supplementary Table S1:**
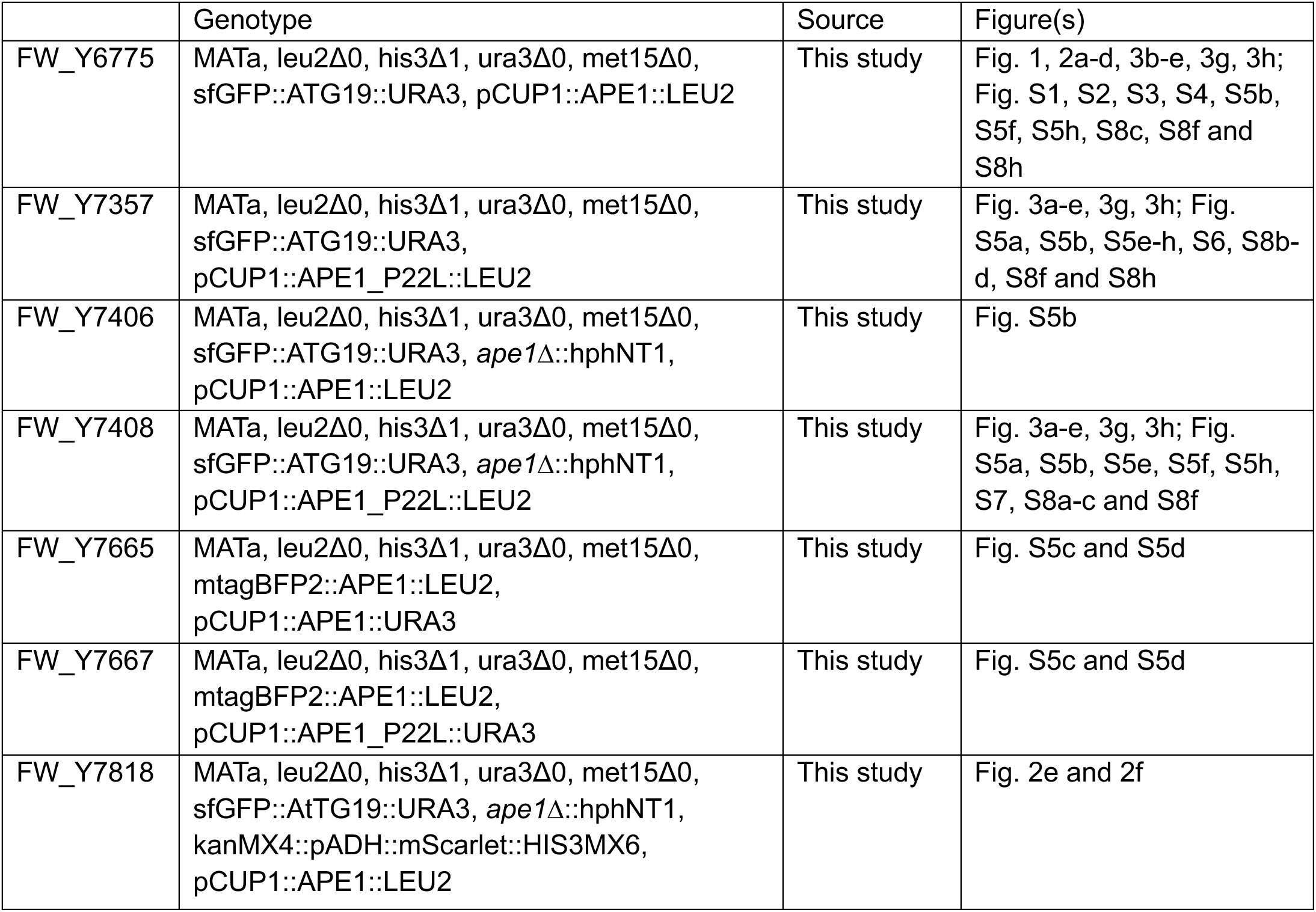
*S. cerevisiae* strains used in this study

**Supplementary Table S2:** Plasmids used in this study

| Name | Genotype | Source | Figure(s) |
| --- | --- | --- | --- |
| FWP_853 | pRS315-pCUP1-Ape1 | Claudine Kraft | Fig. 1, 2, 3b-e, 3g, 3h; Fig. S1, S2, S3, S4, S5b, S5f, S5h, S8c, S8f and S8h |
| FWP_949 | pRS315-pCUP1-Ape1(P22L) | This study | Fig. 3a-e, 3g, 3h; Fig. S5a, S5b, S5e-h, S6, S7, S8a-d, S8f and S8h |
| FWP_982 | yCplac33-pCUP1-Ape1 | This study | Fig. S5c and S5d |
| FWP_1035 | yCplac33-pCUP1-Ape1(P22L) | This study | Fig. S5c and S5d |
| pHM220 | T7_StrepII-NTtag-HRV3C-Ape1_T7>pET21a | This study | Fig. 5c and 5d; Fig. S10h-k and S10m |
| pHM221 | T7_StrepII-NTtag-HRV3C-Ape(P22L)_T7>pET21a | This study | Fig. 5c, 5d; Fig. S10h-j, S10l and S10m |

**Supplementary Table S3:** WT and P22L Ape1 condensate cryo-ET data acquisition parameters

|  |  |
| --- | --- |
| Microscope | Titan Krios G4 |
| Camera | Thermo Scientific Falcon 4i |
| Energy filter | Selectris X |
| Acquisition software | SerialEM |
| Accelerating voltage (kV) | 300 |
| Nominal magnification | 64000 |
| Nominal pixel size (Å/px) | 1.971 |
| Total electron dose (e<sup>-</sup>/Å<sup>2</sup>) | ~150 |
| Acquisition scheme | Dose-symmetric with initial stage tilt of ±8° |
| Tilt increment (°) | 2 |
| Tilt range (°) | ±60 from initial stage tilt |
| Target defocus range (µm) | -1.5 to -4.5 |
| Energy filter width | 10eV |

**Supplementary Table S4:** WT and P22L Ape1 subtomogram averaging summary

|  | WT | P22L with endogenous WT | P22L without endogenous WT |
| --- | --- | --- | --- |
| Number of tomograms acquired | 30 | 8 | 19 |
| Number of tomograms used for template matching | 9 | 6 | 6 |
| Final number of particles | 12370 | 20518 | 6892 |
| Final resolution (Å) | 3.84 | 5.6 | 9.1 |

**Supplementary Table S5:**
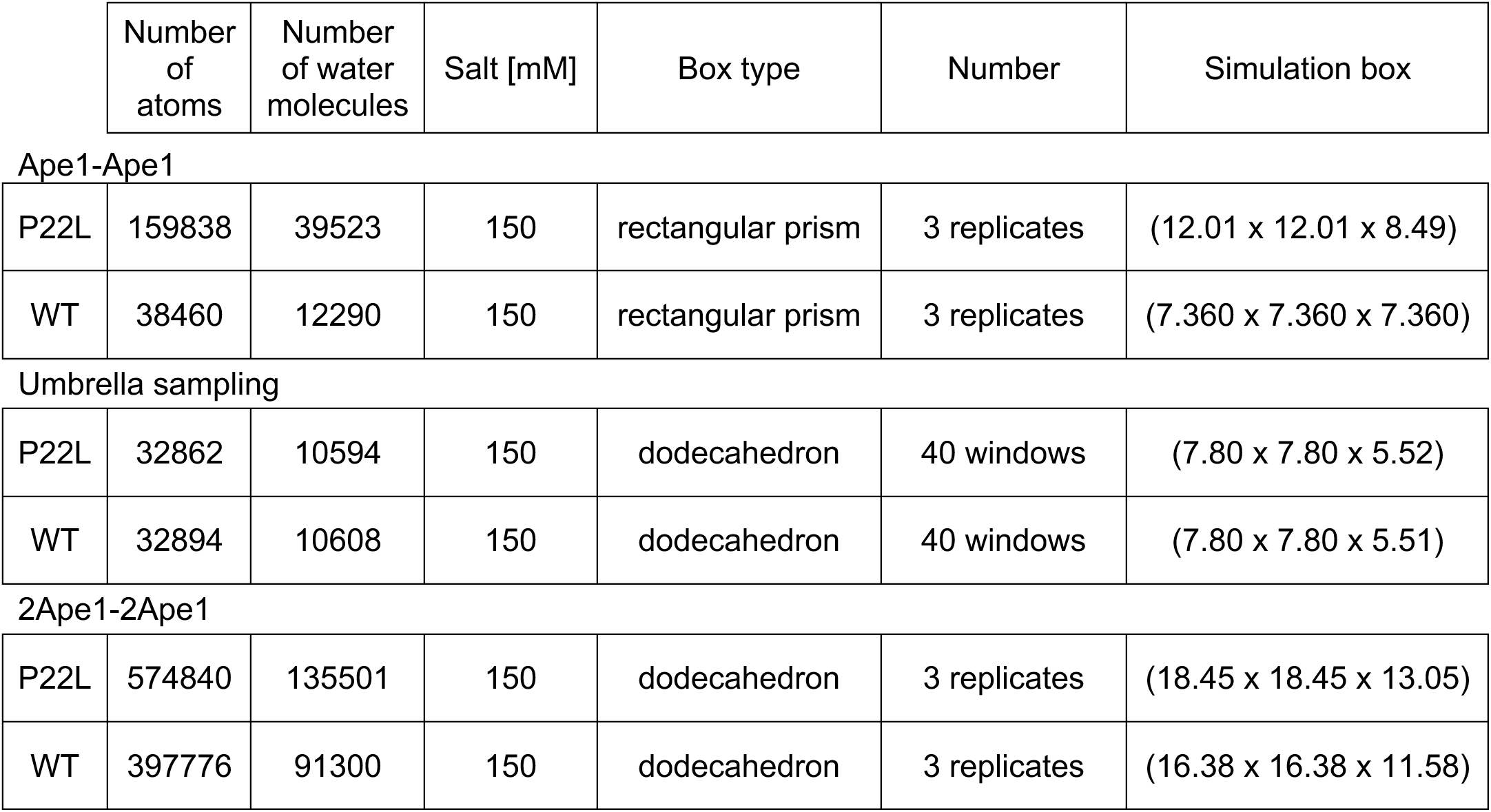
Simulation parameters summary

|  | Number of atoms | Number of water molecules | Salt [mM] | Box type | Number | Simulation box |
| --- | --- | --- | --- | --- | --- | --- |
| **Ape1-Ape1** |  |  |  |  |  |  |
| P22L | 159838 | 39523 | 150 | rectangular prism | 3 replicates | (12.01 x 12.01 x 8.49) |
| WT | 38460 | 12290 | 150 | rectangular prism | 3 replicates | (7.360 x 7.360 x 7.360) |
| **Umbrella sampling** |  |  |  |  |  |  |
| P22L | 32862 | 10594 | 150 | dodecahedron | 40 windows | (7.80 x 7.80 x 5.52) |
| WT | 32894 | 10608 | 150 | dodecahedron | 40 windows | (7.80 x 7.80 x 5.51) |
| **2Ape1-2Ape1** |  |  |  |  |  |  |
| P22L | 574840 | 135501 | 150 | dodecahedron | 3 replicates | (18.45 x 18.45 x 13.05) |
| WT | 397776 | 91300 | 150 | dodecahedron | 3 replicates | (16.38 x 16.38 x 11.58) |

### Supplementary Video Legends

**Supplementary Video S1: Representative example of a denoised WT Ape1 tomogram reconstruction.** After scrolling through the tomogram volume in one direction, low-pass filtered electron density maps of the Ape1 dodecamer are placed back at particle positions identified by template matching and refined during the subtomogram averaging. Manually refined segmented ER membranes are also shown. A 90° rotation of the segmentation about the x axis reveals the morphology of the cap of the Ape1 complex captured in this tomogram.

**Supplementary Video S2: All atom molecular dynamics simulation of two WT Ape1 dimers representing the edges of two neighbouring Ape1 dodecamers.** The propeptides interact via their N-terminal helices as an antiparallel 4-helix bundle, as predicted by AlphaFold (Fig. S8j). Additional Ape1 monomers shown in grey are only included for visualisation and were not part of the simulation. Movement of the two Ape1 dimers relative to each other highlights the flexibility of the propeptide C-terminus upon propeptide-propeptide interaction.

**Supplementary Video S3: Coarse-grained molecular dynamics simulation to recapitulate the dynamics of the WT Ape1 condensate.** Patchy particles represent Ape1 dodecamers, as shown in Fig. 5a and the interaction strength approximates the WT propeptide-propeptide interaction strength. Fast particle exchange and formation of spherical condensates is observed during the timecourse of the simulation.

**Supplementary Video S4: Coarse-grained molecular dynamics simulation to recapitulate the dynamics of the P22L Ape1 condensate.** Patchy particles represent Ape1 dodecamers, as shown in Fig. 5a and the interaction strength approximates the P22L propeptide-propeptide interaction strength. Slow particle exchange and formation of non-spherical condensates is observed during the timecourse of the simulation.

## Notes

### Competing Interest Statement

The authors have declared no competing interest.

